# Genome mining yields new disease-associated ROMK variants with distinct defects

**DOI:** 10.1101/2023.05.05.539609

**Authors:** Nga H. Nguyen, Srikant Sarangi, Erin M. McChesney, Shaohu Sheng, Aidan W. Porter, Thomas R. Kleyman, Zachary W. Pitluk, Jeffrey L. Brodsky

**Affiliations:** Department of Biological Sciences, A320 Langley Hall, University of Pittsburgh, Pittsburgh, PA, 15260, USA; Paradigm4, Inc., Suite 360, 281 Winter Street, Waltham, MA, 02451, USA; Renal-Electrolyte Division, School of Medicine, S929 Scaife Hall, University of Pittsburgh PA, 15261, USA

## Abstract

Bartter syndrome is a group of rare genetic disorders that compromise kidney function by impairing electrolyte reabsorption. Left untreated, the resulting hyponatremia, hypokalemia, and dehydration can be fatal. Although there is no cure for this disease, specific genes that lead to different Bartter syndrome subtypes have been identified. Bartter syndrome type II specifically arises from mutations in the *KCNJ1* gene, which encodes the renal outer medullary potassium channel, ROMK. To date, over 40 Bartter syndrome-associated mutations in *KCNJ1* have been identified. Yet, their molecular defects are mostly uncharacterized. Nevertheless, a subset of disease-linked mutations compromise ROMK folding in the endoplasmic reticulum (ER), which in turn results in premature degradation via the ER associated degradation (ERAD) pathway. To identify uncharacterized human variants that might similarly lead to premature degradation and thus disease, we mined three genomic databases. First, phenotypic data in the UK Biobank were analyzed using a recently developed computational platform to identify individuals carrying *KCNJ1* variants with clinical features consistent with Bartter syndrome type II. In parallel, we examined ROMK genomic data in both the NIH TOPMed and ClinVar databases with the aid of a computational algorithm that predicts protein misfolding and disease severity. Subsequent phenotypic studies using a high throughput yeast screen to assess ROMK function—and analyses of ROMK biogenesis in yeast and human cells—identified four previously uncharacterized mutations. Among these, one mutation uncovered from the two parallel approaches (G228E) destabilized ROMK and targeted it for ERAD, resulting in reduced protein expression at the cell surface. Another ERAD-targeted ROMK mutant (L320P) was found in only one of the screens. In contrast, another mutation (T300R) was ERAD-resistant, but defects in ROMK activity were apparent after expression and two-electrode voltage clamp measurements in *Xenopus* oocytes. Together, our results outline a new computational and experimental pipeline that can be applied to identify disease-associated alleles linked to a range of other potassium channels, and further our understanding of the ROMK structure-function relationship that may aid future therapeutic strategies.

**Author Summary:** Bartter syndrome is a rare genetic disorder characterized by defective renal electrolyte handing, leading to debilitating symptoms and, in some patients, death in infancy. Currently, there is no cure for this disease. Bartter syndrome is divided into five types based on the causative gene. Bartter syndrome type II results from genetic variants in the gene encoding the ROMK protein, which is expressed in the kidney and assists in regulating sodium, potassium, and water homeostasis. Prior work established that some disease-associated ROMK mutants misfold and are destroyed soon after their synthesis in the endoplasmic reticulum (ER). Because a growing number of drugs have been identified that correct defective protein folding, we wished to identify an expanded cohort of similarly misshapen and unstable disease-associated ROMK variants. To this end, we developed a pipeline that employs computational analyses of human genome databases with genetic and biochemical assays. Next, we both confirmed the identity of known variants and uncovered previously uncharacterized ROMK variants associated with Bartter syndrome type II. Further analyses indicated that select mutants are targeted for ER-associated degradation, while another mutant compromises ROMK function. This work sets-the-stage for continued mining for ROMK loss of function alleles as well as other potassium channels, and positions select Bartter syndrome mutations for correction using emerging pharmaceuticals.

## Introduction

First identified in 1962, Bartter syndrome is group of rare, life-threatening disorders caused by defects in or impaired function of electrolyte channels within the kidney, compromising renal sodium and potassium handling and resulting in excessive electrolyte and water excretion (1). To date, therapies for Bartter syndrome include electrolyte supplements and non-steroidal anti-inflammatory drugs, but are limited to mitigating the symptoms, such as electrolyte supplementation and the use of non-steroidal anti-inflammatories. Although disease severity, presentation, and age of onset vary, Bartter syndrome can lead to a failure to thrive, sudden cardiac arrest, and even death (2, 3).

One among several causes of Bartter syndrome arises from defects in a potassium channel residing on the apical surface of two segments of the nephron: the thick ascending limb and the cortical collecting duct (4). The channel, ROMK (also known as Kir1.1), is encoded by *KCNJ1* and was the first inward rectifying potassium (Kir) channel identified (5–7). Like other Kir channels, ROMK functions as a tetramer (8) and exhibits a larger inward current than outward current; all family members also share a common structure that contains two transmembrane domains (TMD) and cytoplasmic N- and C-terminal domains (9). In the kidney, ROMK plays a central role in mediating potassium efflux, which in turn provides a crucial source of potassium to facilitate sodium reabsorption through the sodium potassium-chloride cotransporter (NKCC2) in the thick ascending limb. Furthermore, ROMK-dependent potassium secretion in the thick ascending limb generates a lumen positive transepithelial potential that drives paracellular sodium absorption (10). Mutations in ROMK specifically give rise to Bartter syndrome type II, also called antenatal Bartter syndrome, since patients often present prenatally (e.g., with excessive amniotic fluid). Among these individuals, observed features include a failure to thrive, renal salt wasting and volume depletion, early post-natal hyperkalemia, hypercalcuria, nephrocalcinosis, and arrhythmias, all of which contribute to a high infant mortality rate (11).

In theory, defects in ROMK might arise from a lack of expression, defects in protein folding and/or tetramerization, the accelerated degradation of poorly folded/assembled subunits, inefficient transport to the cell surface, and/or altered channel (i.e., potassium transport) activity. Indeed, early studies in *Xenopus* oocytes and COS-7 cells demonstrated that some Bartter syndrome type II-associated mutants were absent from the cell surface and others were defective for potassium transport (12–14). Later work by our group showed that four disease-causing mutations in ROMK that cluster in a cytosolic, β sheet-rich immunoglobulin domain cause the protein to misfold in the endoplasmic reticulum (ER) (15), an outcome that targets ROMK for ER associated degradation, or ERAD.

The ERAD pathway represents a first-line defense in the secretory pathway to recognize and deliver misfolded proteins to the ubiquitin-proteasome system (UPS), which resides on the cytosolic face of the ER. During ERAD, molecular chaperones, such as heat shock protein 70 (Hsp70), recognize and target misfolded proteins both for extraction (or “retrotranslocation”) from the ER lumen and ER membrane into the cytosol and ubiquitination, which serves as a prelude to proteasome-dependent degradation (16–21). Retrotranslocation requires a AAA+-ATPase, known as Cdc48 in yeast, or p97 (also known as Valosin Containing Protein; VCP) in higher cells (22, 23). In a study utilizing both a new yeast expression system and human cell lines, we showed that Hsp70 and Cdc48 were required for the degradation of Bartter syndrome-linked mutant ROMK species, whereas wild-type ROMK was relatively stable (15). In addition, the expression of ROMK in a yeast strain lacking two endogenous potassium channels (*trk1*Δ*trk2*Δ) restored growth on low potassium media (24, 25). As a result, ROMK folding, trafficking to the plasma membrane (where it functions), and potassium transport can be assayed in yeast. Together, these data indicate that the yeast system can effectively monitor the efficacy of ROMK biogenesis and provides a facile growth assay, allowing one to screen for defective ROMK mutations in a quantitative and high-throughput manner.

The rapid growth of human genome sequence data and improved curation of existing databases have facilitated the identification of disease-linked genes as well as uncharacterized disease-causing mutations. To date, ROMK mutations associated with Bartter syndrome type II were primarily identified via clinical studies (4, 12, 26, 27), but numerous uncharacterized disease-linked ROMK mutations likely remain unearthed in human databases.

To identify and characterize some of these mutations, we now report on the use of two computational approaches to uncover additional ROMK variants associated with Bartter syndrome type II. First, we examined ROMK missense mutations in two NIH-supported databases, the Trans-Omics for Precision Medicine (TOPMed) database (28) and ClinVar (29), using an algorithm that predicts mutation severity. The algorithm, known as Rhapsody (30), utilizes evolutionary conservation along with structural and dynamic features. We previously validated Rhapsody’s predictive power to probe the potential impact of a small group of known ROMK mutations (31). Second, we performed *in silico* association analyses to identify links between ROMK variants in the UK Biobank and disease-associated phenotypes (32–34) using the REVEAL: Biobank computational platform (35–38). As a result of these complementary approaches, we report here on the identification and characterization of a cohort of ROMK variants using yeast, *Xenopus* oocytes, and tissue culture cells. Ultimately, we discovered new variants that are unstable and targeted for ERAD, those that are poorly expressed at the cell surface, and/or those that exhibit defective channel function. The identification of a common allele from the two approaches validated the complementary nature of these approaches and outlines a new pipeline to assess other identified disease-associated mutations in ROMK, an effort that may aid in the development of precision medicine to treat those with Bartter syndrome type II.

## Results

### A computationally-guided analysis reveals uncharacterized ROMK mutations associated with Bartter Syndrome type II

To isolate previously uncharacterized ROMK mutations associated with Bartter syndrome type II, we first analyzed genomic data collected from the TOPMed program. TOPMed is an NIH-sponsored whole genome sequencing program with a cohort of more than 180,000 participants who have lung, heart, sleep, and blood disorders (28, 39). Moreover, genomic data from the cohort are continuously deposited into the publicly available Bravo browser (39). At the time of our analysis, a total of 758 ROMK variants were observed in 128,568 individuals, 124 of which are missense mutations (see **S1 Table**). To assess the potential disease severity of each of these amino acid substitutions, we then used Rhapsody, a computational algorithm that was first developed to analyze amino acid variants based on sequence conservation, structure, dynamics, and coevolutionary features (30). We previously verified the power of this *in silico* method to correctly assess the impact of a randomly selected series of ROMK mutations on yeast growth in an assay that reports on ROMK plasma membrane residence and channel function (see **Introduction**) (31). We also found that Rhapsody predicted the severity of known disease-linked ROMK mutations with >90% accuracy.

Along with this computational analysis, we examined potential disease association amongst the 124 TOPMed mutations by cross-examining the NIH ClinVar database, a public archive of human genomic variants and their evidence-based clinical interpretations (29). Ultimately, we focused on mutations located in regions required for protein function and folding, e.g., the PIP2-binding domain (40) and the immunoglobulin-like fold, as well as mutations designated as having “uncertain clinical significance” for Bartter syndrome in ClinVar (29) (**Fig 1A**).

**Figure 1.**
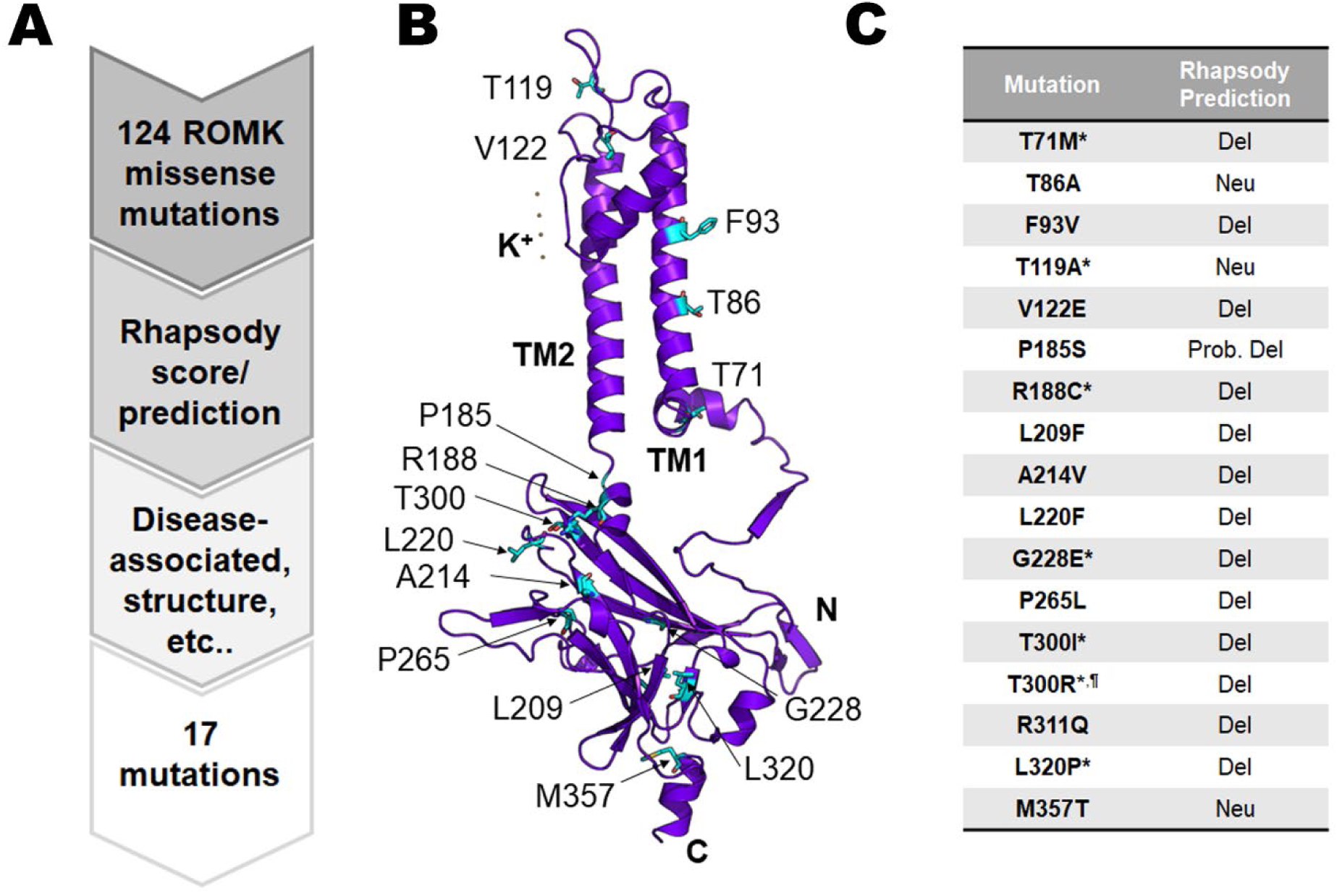
Computer-guided analysis of TOPMed and ClinVar databases to identify previously uncharacterized ROMK missense mutations associated with Bartter syndrome type II. (A) A flowchart describing how 17 mutations were selected for further examination. All missense ROMK mutations available in the Bravo database (39), in which data from the TOPMed study are continuously deposited, were analyzed. At the time of this study, the Bravo database was in its “freeze 5” version, where a total of 124 missense ROMK mutations were available. We then analyzed all 124 mutations using the computational algorithm Rhapsody (see text for details) to predict mutation pathogenicity, and picked 16 mutations based on Rhapsody score, disease (Bartter syndrome) association, and location in the ROMK structure. One additional mutation from the ClinVar database (T300R) was chosen because it resided in the same residue as a TOPMed mutation (T300I). (B) Location of the 17 mutations on a ROMK homology model. While ROMK tetramerizes to form a functional channel, only one monomer is shown. Seventeen residues of interest are shown in light blue sticks. The homology model was built based on the crystal structure of Kir2.2 (PDB ID: 3SPG), which is 47.42% identical to ROMK1. Images were rendered using PyMOL (ver. 2.6.0). (C) List of 17 mutations and their Rhapsody predictions. “Del” = deleterious, “Neu” = neutral, or predicted to have no effects on channel architecture. A designation of “Prob. Del” indicates that the Rhapsody score is close to the 0.5 cutoff. For example, the Rhapsody score of P185S = 0.549 and is thus listed as “Prob. Del”. * denotes an uncharacterized Bartter mutation, which is defined as a disease-associated mutation in ClinVar, but is listed as having uncertain clinical significance, ¶ denotes the mutation obtained solely from ClinVar. A comprehensive table of Rhapsody scores and prediction for the 17 mutations of interest, as well as all 124 TOPMed mutations, can be found in **S1** and **S2 Tables**, respectively.

Based on these analyses, representative mutations were chosen for further assessment (**Fig 1** and **S2 Table**). Most of the mutations (12 out of 17) reside in the ROMK cytoplasmic domain (**Fig 1B**), which contains key regions that play important roles during protein folding and for channel function. These regions include the cytoplasmic pore, the G-loop, and the PIP_2_-binding pocket. For example, T300I is located on a β sheet proximal to the G-loop region, which is solvent-accessible at the top of the cytoplasmic pore and regulates channel gating and inward rectification in ROMK and other Kir channels (41–43). Given the potential contribution of the T300 site to channel function, we also added T300R for further analysis, which is a residue that was identified in ClinVar. Ultimately, amongst the 17 variants, 14 were predicted to be deleterious, i.e., assigned a Rhapsody score of ≥ 0.5, with G228E having the highest Rhapsody score (0.930; **Fig 1C** and **S2 Table**). Interestingly, G228E resides in the β sheet-rich immunoglobulin domain, where—as noted above—mutations have been shown to compromise ROMK folding and stability (15). It is also noteworthy that the mutation with the highest frequency in the population (0.68%), M357T, is predicted to be neutral.

ClinVar predicts that seven mutations are linked to Bartter syndrome, though they are classified as having uncertain clinical significance (**Fig 1C**, mutations marked with a *). In contrast, seven other mutations were previously associated with Bartter syndrome and have clear clinical consequences (**S2 Table**, denoted by “Bartter” in the Background information column). For example, T86A is listed on ClinVar as likely being benign, consistent with its neutral Rhapsody score, while another mutation with a deleterious Rhapsody score, P185S, is in the putative PIP_2_-interacting domain and disrupts channel conductance, likely by altering PIP_2_ binding (44). A comprehensive list of the 124 TOPMed variants and the 17 chosen mutations, along with their Rhapsody predictions, can be found in **S1** and **S2 Tables**, respectively.

We next assayed the 17 mutants in the yeast growth assay that assesses both potassium channel residence at the cell surface and function. As noted in the **Introduction**, the basis of this assay is that yeast lacking two endogenous potassium channels, Trk1 and Trk2, require the presence of a functional exogenous potassium channel at the plasma membrane to support growth on low potassium (**Fig 2A**) (24, 45, 46). We and others have used the corresponding *trk1*Δ*trk2*Δ yeast strain to characterize mutant alleles and the channel properties of ROMK and several other Kir channels, along with members of other potassium channel classes (15, 25, 47–50).

**Figure 2.**
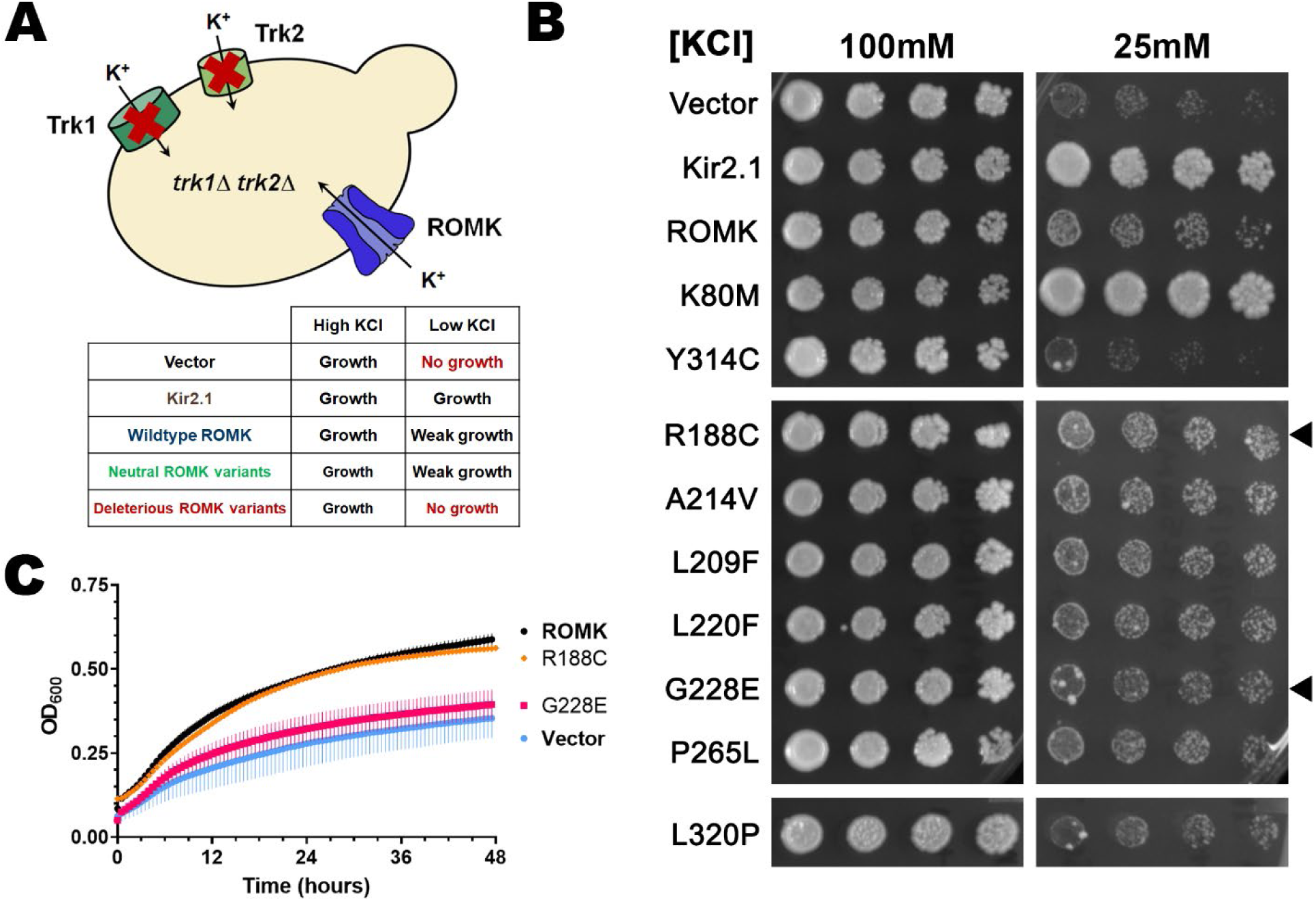
ROMK mutations from TOPMed and ClinVar inserted into a yeast expression vector show varying growth defects on low potassium medium. (A) Schematic of a yeast-based assay to assess the activity of a human potassium channel. A yeast strain lacking endogenous potassium transporters, Trk1 and Trk2, is viable, yet unable to grow on medium containing low potassium, unless a human potassium channel (e.g., Kir2.1 or ROMK) is expressed. Because of impaired ROMK activity at low pH (78) and exaggerated steady-state residence in the ER (48), ROMK exhibits only a weak growth phenotype on low potassium in contrast to Kir2.1. The table shows the expected growth phenotype of yeast containing an empty vector, or expressing a related Kir channel, Kir2.1, or ROMK, and the predicted growth phenotype of yeast expressing a ROMK mutation from Fig 1. (B) Yeast viability assays on solid medium. Yeast cultures were transformed with an empty expression vector as a negative control, or with a plasmid expressing Kir2.1, ROMK, or the indicated ROMK mutation. Kir2.1 and a known hyperactive ROMK mutation, K80M, were used as positive controls, while an unstable Bartter mutant (Y314C) (15, 51) was used as a negative control. Yeast cultures were grown overnight to saturation, diluted the next day, and serial 1-to-5 yeast dilutions were spotted on solid medium containing high (100 mM) or low (25 mM) potassium. Here, the growth phenotype of 7 representative mutants is shown. The black triangles point to ROMK and mutations either with largely no effect on growth (R188C) or with a growth defect on low potassium medium (G228E). (C) Yeast viability assays were performed in liquid medium supplemented with low potassium (25 mM). Yeast containing a vector control, or expressing ROMK, or a ROMK mutant (e.g., R188C and G228E, as shown here) were grown overnight to saturation and diluted the next day to an OD_600_ of 0.20 with medium supplemented with 25 mM KCl. OD_600_ readings were recorded, normalized to wells containing only medium, and OD_600_ readings over the course of 48 hr are shown. Graphs were made using GraphPad Prism (ver. 9.5.0), and data represent results from two replicates, ± S.E. (error bars). Growth assays on solid and in liquid medium of yeast expressing each of the 17 mutations are shown in **S1** and **S2 Figs**, respectively, and **S3 Table** summarizes the results from the growth assays.

The wild-type and each of the 17 mutant ROMK proteins were expressed in the *trk1*Δ*trk2*Δ yeast strain, and growth on solid medium as well as in liquid medium in a 96- well plate assay were measured (**Fig 2B-C** and **S1-S2 Figs**). Yeast containing an empty vector or expressing Y314C, a Bartter mutation previously shown to compromise ROMK folding and function (14, 15, 51), were used as negative controls, while yeast expressing a related Kir channel (Kir2.1) that traffics more efficiently to the cell surface than ROMK (48, 49), along with a known hyperactive ROMK mutant allele (K80M) (52), were used as positive controls. Based on yeast growth in low potassium-containing media, we first noticed that G228E, the mutation in the immunoglobulin fold with the most deleterious Rhapsody score exhibited an expected severe growth defect, i.e., growth was similar to that of yeast containing a vector control (**Fig 2B-C**). These data are consistent with G228 residing in the β sheet-rich immunoglobulin-like domain (see above and (51)); other disease-associated mutations in this region, including our negative control, Y314C, are rapidly targeted for ERAD, likely due to severe folding defects (15). Another mutation located at the base of the immunoglobulin-like β sheet-containing sandwich domain, L320P, similarly prevented yeast growth on solid medium (**Fig 2B**), but only modestly impaired growth in liquid medium (**S2 Fig**). These data may reflect the fact that this mutation has a Rhapsody score (0.588) that predicts a marginal defect. Perhaps the magnitude of the defect rises when L320P must support growth under more anoxic conditions (i.e., in solid medium) in contrast to growth in aerated cultures (i.e., in liquid medium). Based on their growth phenotypes in low potassium, we classified the 17 mutants into four groups based on their growth phenotypes: a severe defect (e.g., G228E), a moderate defect (e.g., L320P), a slight defect, and no defect. **S3 Table** summarizes the growth phenotype of each mutation according to this classification.

Importantly, some mutations exhibited growth phenotypes in accordance with previous work and with their Rhapsody scores. One such example is the L220F Bartter mutation, which grew more slowly on solid and in liquid media. These data are consistent with a highly deleterious Rhapsody score (0.759), a “pathogenic” classification in ClinVar (**S1 Table**), and previous studies demonstrating reduced channel currents in *Xenopus* oocytes (14, 26). The defect in channel function caused by this mutation likely stems from its localization adjacent to S219, a phosphorylation site for protein kinase A (53), which is crucial for maintaining the open state of the channel (54). In addition, yeast expressing the likely benign and predicted neutral M357T variant grew robustly, as anticipated. Besides M357T, yeast expressing the two other predicted neutral mutations similarly showed minimal or no growth defects (T86A and T119A, see **S1-2 Figs** and **S3 Table**). Yet, the concordance between Rhapsody predictions and growth phenotypes was not absolute. A Bartter mutation with a “probably deleterious” designation and reported to likely affect PIP_2_ binding [P185S; (44)] was also without consequence in yeast. However, this mutation reduced single channel conductance only when PIP_2_ was depleted, and had no apparent effect on channel currents or surface expression in *Xenopus* oocytes (44), which might explain the lack of a yeast growth defect.

Ultimately, we selected four alleles to characterize in greater depth. G228E and L320P were chosen for their respective strong (G228E) and more moderate (L320P) growth defects in yeast viability assays and for their status of having clinically uncertain significance based on ClinVar. We also selected T86A as a representative neutral mutation that exhibited no growth defect, and T300R due to its moderate growth defect and the importance of the G-loop in supporting ROMK function/stability, as described above.

### Phenotype-guided association analyses of the UK Biobank identified additional disease-associated ROMK variants

In parallel to the data mining protocol above, we pursued an alternate and more direct strategy to identify previously ill-characterized and novel disease-linked mutations. More specifically, we wished to identify individuals who exhibit features characteristic of Bartter syndrome type II, but are undiagnosed or harbor previously unidentified mutations in *KCNJ1*. Therefore, we utilized the UK Biobank, a genomic and metabolomic resource for multi-omics data retrieved from an ongoing participant study initiated in 2006 (32). In particular, we performed three genome-wide association studies (GWAS) between phenotypic data and ROMK mutations using REVEAL: Biobank, an analytical platform built upon SciDb (55) that supports elastic scaling for efficient and cost-effective genomic analyses (35–38) (**S3 Fig**). We utilized whole exome sequencing data, which at the time of this study, had been made available for ∼200,000 UK Biobank participants (56, 57). Whole exome sequencing measures the coding regions of the genome and helps identify disease-causing and/or rare genetic variants. Combined with the large sample size of the UK Biobank cohort and rich phenotypic datasets, whole exome sequencing can also help elucidate gene function, which is otherwise challenging with imputed genomic data (58–61) and may require the application of additional statistical methods to compensate for missing data (62). Within this whole exome sequencing dataset, there were 511 *KCNJ1* variants (**S4 Table**), and after applying a minor allele frequency (maf) filter (maf > 1e-05), we selected 142 variants for the GWAS analyses.

The first association study employed 15 continuous phenotypes for their relevance to ROMK function, to Bartter syndrome type II, and to hypertension (9, 63), and included phenotypes such as systolic and diastolic blood pressure, serum urea, creatinine, calcium, and phosphate, as well as urine potassium and sodium. For the second analysis, we selected 15 unique phenotypic codes, or “Phecodes”, associated with Bartter syndrome type II. The use of Phecodes has recently emerged as an effective route to classify clinical phenotypes and is thus suited to phenome-wide association studies compared to traditional billing ICD10 codes (34). For example, the 15 Phecodes we selected represent 25 traditional ICD10 codes. The Phecodes chosen include clinical Bartter syndrome phenotypes related to fatigue and weakness (e.g., malaise and fatigue), volume loss and thirst (e.g., polyuria), hyperaldosteronism, electrolyte imbalance, and neonatal diagnoses. Notably, the analysis excluded Phecodes linked to hypothyroidism and diabetes. We reasoned that these phenotypes might increase the number of false negatives since they are not directly related to ROMK. Finally, we used available metabolomics data from the Biobank for the third association study (**Fig 3A**).

**Figure 3.**
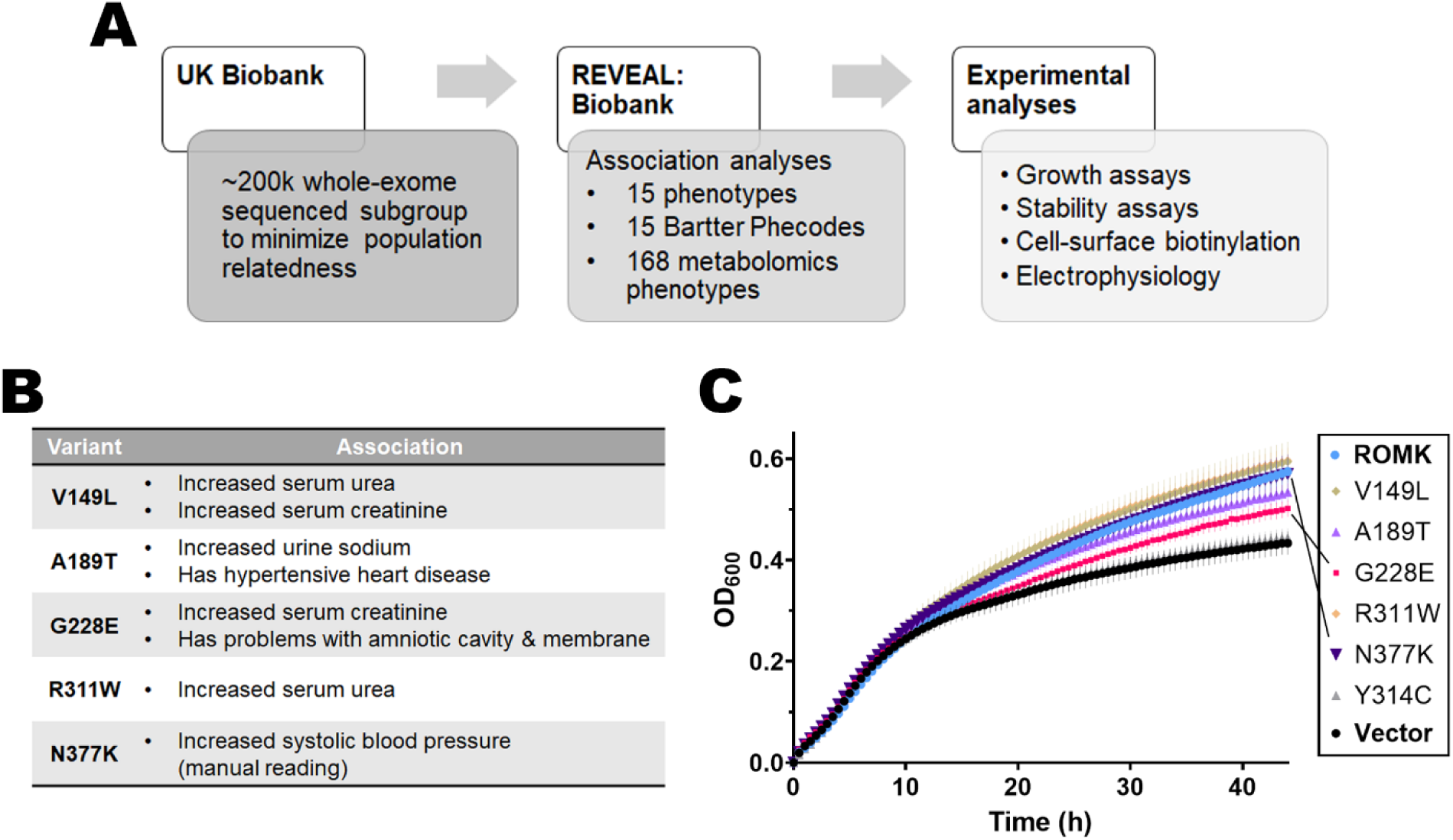
Mining the UK Biobank to identify ROMK mutations associated with disease-related phenotypes and show growth defects in yeast. (A) A flowchart describing how the UK Biobank was mined to search for potential disease-causing mutations. From a sub-population of the UK Biobank (see text for details) that contains genomic and phenotypic data from ∼200k participants, we performed *in silico* analysis to find significant associations between ROMK variants and disease-related phenotypes. REVEAL: Biobank, a computational platform built upon SciDB and featuring elastic scaling was used to analyze data (See **S3 Fig** for details). Three analyses were performed: disease-associated phenotypes (Bartter syndrome and blood pressure-related), disease Phecodes, and metabolomic phenotypes, and a select group of five mutants was chosen for further experimental analysis. (B) Table summarizing the list of potential disease-related ROMK mutations and their associated phenotypes. For a more comprehensive results of the phenotypic association analysis, see **Tables 1-3** and **S5 Table**. (C) Graph shows yeast viability assays in liquid medium supplemented with low potassium (25 mM). Yeast containing a vector control, or expressing ROMK, or a ROMK mutant corresponding to a variant from the UK Biobank were grown overnight to saturation and diluted the next day to an OD_600_ of 0.20 with medium supplemented with 25 mM KCl. OD_600_ readings were recorded, normalized to wells containing only medium and the first time point, and OD_600_ readings over the course of 44 hr are shown. Graphs were made using GraphPad Prism (ver. 9.5.0), and data represent results from ten replicates, ± S.E. (error bars).

**Table 1.** Top significant associations between relevant binary disease phenotypes and ROMK variants in the UK Biobank. Table shows significant associations obtained from GWAS performed with two algorithms, SAIGE (64) and REGENIE (65). which provide similar results with slightly different p-values. Relevant disease phenotypes were selected from a list of Phecodes, i.e., refined groups of International Classification of Diseases (ICD) codes that are both clinically meaningful and facilitate genome analysis (34, 136).

| SAIGE |  |  |  |  |  |  |  |
| --- | --- | --- | --- | --- | --- | --- | --- |
| Phenotype<br>(Phecode #) | Chromosome<br>position | Nucleotide<br>change | P-value | Observation<br># | Consequence | Beta<br>value | Muta-<br>tion |
| Problems associated with<br>amniotic cavity and<br>membranes (653.00) | 11:128839618 | C > T | 6.07E<br>-04 | 122586 | Missense | 22.6 | G228E |
| Hypertensive Heart<br>Disease (401.21) | 11:128839736 | C > T | 7.26E<br>-04 | 107442 | Missense | 222.955 | A189T |
| REGENIE |  |  |  |  |  |  |  |
| Phenotype<br>(Phecode #) | Chromosome<br>position | Nucleotide<br>change | P-value | Observation<br># | Consequence | Beta<br>value | Muta-<br>tion |
| Hypopotassemia (276.14) | 11:128839710 | G > A | 7.84E<br>-04 | 121659 | Synonymous | 5.0 | N197N |
| Electrolyte imbalance<br>(276.10) | 11:128839710 | G > A | 9.26E<br>-04 | 121692 | Synonymous | 4.8 | N197N |

**Table 3.** Comparison of metabolite levels between wild-type versus individuals carrying ROMK variants. Disease phenotypes with significant associations with mutations in *KCNJ1* from **Table 2** are listed in the first column. “#” denotes the number of individuals with the mutation or the wild-type allele, and data shown are the means, medians, and standard deviations (“S.D.”) of each phenotype.

| Phenotype | Chromosome position | Units | Mutation | Individuals with mutation |  |  |  | Wildtype individuals |  |  |  |
| --- | --- | --- | --- | --- | --- | --- | --- | --- | --- | --- | --- |
|  |  |  |  | # | Mean | Median | S.D. | # | Mean | Median | S.D. |
| Urea | 11:128839052 | mmol/L | 3' UTR | 36744 | 5.45 | 5.32 | 1.38 | 93583 | 5.42 | 5.28 | 1.38 |
| Phosphate | 11:128839052 | mmol/L | 3' UTR | 33846 | 1.16 | 1.16 | 0.16 | 85997 | 1.17 | 1.17 | 0.16 |
| Creatinine | 11:128839856 | mmol/L | V149L | 2 | 0.1 | 0.1 | 0.004 | 32081 | 0.07 | 0.06 | 0.01 |
| Urea | 11:128839856 | mmol/L | V149L | 12 | 6.49 | 6.78 | 1.15 | 131199 | 5.43 | 5.29 | 1.38 |
| Urea | 11:128839370 | mmol/L | R311W | 15 | 6.62 | 6.78 | 1.04 | 131197 | 5.43 | 5.29 | 1.38 |
| Creatinine (enzymatic) in urine | 11:128839618 | umol/L | G228E | 23 | 11863.52 | 12760 | 6323.81 | 133898 | 8724.24 | 7410 | 5680.85 |
| Sodium in urine | 11:128839736 | mmol/L | A189T | 4 | 129.6 | 131.75 | 35.63 | 133636 | 75.22 | 66.4 | 42.96 |
| Systolic blood pressure (manual) | 11:128839170 | mmHg | N377K | 4 | 173 | 168.5 | 23.85 | 7800 | 140.67 | 139 | 19.89 |
| Systolic blood pressure (automated) | 11:128839170 | mmHg | N377K | 27 | 132.74 | 135 | 16.46 | 129973 | 140.07 | 139 | 19.55 |

SAIGE (64) and REGENIE (65) were the two algorithms utilized to perform the GWAS analyses. Both algorithms are standards in bioinformatics workflows and identify significant associations, which represent the computed p-value of a regression test between a mutation and a phenotype in the population tested (see **Materials and Methods**). Results obtained from both algorithms were similar, albeit with minor differences observed in the strength of the association as measured by the p-value. The top significant associations identified between *KCNJ1* variants and Bartter-syndrome relevant phenotypes are presented in **Table 1** (for binary Phecodes) and **Table 2** (for quantitative/ continuous and metabolomic phenotypes). Details for each association are presented in each row of the tables and includes names of the phenotype/phecode, chromosomal position, base pair change, p-value of the association, number of participants used for the computation, consequence of the mutation, beta value (i.e., effect size), and amino acid change (if applicable). For each association in **Table 2**, we computed the mean, median, and standard deviation of the phenotype in individuals carrying a mutation (homozygous or heterozygous) versus wild-type individuals. These data are presented in **Table 3**.

**Table 2.** Top significant associations between relevant disease phenotypes and ROMK variants in the UK Biobank. Table shows significant associations obtained from GWAS with two algorithms: SAIGE (64) for the initial analysis, and REGENIE (65) for result confirmation. Fifteen relevant and continuous disease phenotypes were selected from the phenotype list available in the UK Biobank (see **Materials and Methods** for details).

| SAIGE |  |  |  |  |  |  |  |
| --- | --- | --- | --- | --- | --- | --- | --- |
| Phenotype | Chromosome position | Nucleotide change | P-value | Observation # | Consequence | Beta value | Mutation |
| Urea | 11:128839052 | T > A | 3.86E-05 | 131212 | 3' UTR | 0.02 | - |
| Phosphate | 11:128839052 | T > A | 5.11E-04 | 120657 | 3' UTR | -0.87 | - |
| Creatinine | 11:128839856 | C > A | 5.95E-05 | 32083 | Missense | 2.48 | V149L |
| Urea | 11:128839370 | G > A | 2.93E-04 | 131212 | Missense | 0.88 | R311W |
| Sodium in urine | 11:128839736 | C > T | 6.10E-03 | 133640 | Missense | 1.31 | A189T |
| Creatinine (enzymatic) in urine | 11:128839618 | C > T | 9.79E-03 | 133921 | Missense | 0.50 | G228E |
| Systolic blood pressure (automated) | 11:128839170 | G > T | 6.48E-02 | 130002 | Synonymous | -0.33 | N377K |
| Systolic blood pressure (manual) | 11:128839170 | G > T | 9.18E-03 | 7804 | Synonymous | 1.21 | N377K |
| REGENIE |  |  |  |  |  |  |  |
| Phenotype | Chromosome position | Nucleotide change | P-value | Observation # | Consequence | Beta value | Mutation |
| Urea | 11:128839856 | C > A | 9.41E-04 | 131211 | Missense | 0.90 | V149L |

Interestingly, some of the most significant associations (urea and phosphate, **Table 2**, top two rows) were linked to a variant in the 3’ UTR of *KCNJ1*. Future efforts will seek to assay whether this variant affects message expression, stability, translation, and/or targeting. Here, we instead focused on phenotypes with significant associations with missense mutations residing in one of the five *KCNJ1* exons, as summarized in **Fig 3B**. Intriguingly, G228E (**Fig 3B**, **Table 1**, and also see above) once again emerged as an uncharacterized, but likely Bartter-associated, mutation, since individuals carrying this mutation exhibited increased serum creatinine and problems with the amniotic cavity and membrane, both of which manifest in those with Bartter syndrome type II (66). This result supports the power of using complementary computational methods and the predictive power of each protocol.

Other mutations from this analysis were also associated with typical Bartter syndrome phenotypes, such as V149L (increased serum urea and creatinine), A189T (increased urine sodium), and R311W (increased serum urea) (**Fig 3B** and **Table 2**; for details on the means of all metabolite phenotypes associated with each mutation, refer to **Table 3**). Notably, another variant at position R311, R311Q, is a known Bartter mutation and was chosen as one of the 17 TOPMed mutations (**Fig 1** and **S1 Table**). As expected, R311Q expression in *trk1*Δ*trk2*Δ yeast led to slight to moderate growth defects in low potassium media (**S2 Fig** and **S3 Table**). Mutations at this residue disrupt inter-subunit salt bridges and pH-dependent gating (67, 68), which may explain why ROMK channel function is impaired and Bartter syndrome type II arises (14, 69). Finally, we found that heterozygous carriers of N377K had elevated manual systolic blood pressure (**Fig 3B** and **Table 2**), a phenotype atypical of Bartter syndrome in which urinary sodium losses characteristically lead to low blood pressure (70). This might be attributed to the fact that blood pressure is a complex and multi-genotypic trait (71), or possibly due to error in blood pressure measurements (72, 73). Nevertheless, hypertension has been observed in a clinical case study in which a newborn presenting with classical antenatal Bartter syndrome phenotypes (i.e., renal salt wasting and hyperkalemia) also had transient high blood pressure (74). Subsequent genetic testing revealed that the infant carried two mutations in the *KCNJ1* gene, E151K and a deletion of amino acids 116-119, which again strongly supported a diagnosis of antenatal Bartter syndrome. Moreover, the N377K mutation was also identified in the TOPMed program (**S1 Table**) and was therefore chosen for further analysis.

It is worth noting that all individuals with these five mutations (V149L, A189T, G228E, R311W, and N377K) are heterozygous carriers (**S5 Table**), which could contribute to minimal or undiagnosed disease. In addition, the mutations are rare, with each occurring fewer than 27 times in the 200,000-person cohort (**S5 Table**). In contrast, the 3’ UTR variant—see above—has more than 30,000 occurrences, but given its prevalence, the phenotypes linked to this variant could arise from ROMK-independent polymorphisms.

Next, we measured the growth of yeast expressing each variant in liquid medium to begin to assay for functional defects (**Fig 3C**). G228E again showed a growth defect in *trk1*Δ*trk2*Δ yeast on low potassium, as above, consistent with its highly deleterious Rhapsody score. Another predicted-deleterious mutation at the base of the transmembrane domains, A189T (Rhapsody score 0.644), similarly caused a yeast growth defect, albeit to a lesser extent. Meanwhile, V149L, a mutation in the extracellular domain that organizes the potassium selectivity filter, grew as well as the wild-type control, perhaps reflecting its low Rhapsody score (0.193). In contrast, robust growth of yeast expressing the remaining two mutations (R311W and N377K) was observed in low potassium-containing media. The Rhapsody scores for these two mutations are 0.781 and 0.287, respectively. (Note that the structure at N377 is absent from the homology model, which only covers residues 38-364, so an independent Rhapsody analysis of this mutation was performed using a predicted ROMK monomeric model obtained from AlphaFold (75); also see **Discussion**). In contrast, the lack of a growth defect in yeast expressing R311W was surprising since previous studies in *Xenopus* oocytes showed that this mutation reduced channel currents (14, 69). Perhaps the discrepancy can be attributed to the difference in intracellular pH in the two systems, which is pH 4-5 in yeast (76) and pH ∼7.5 in *Xenopus* oocytes (77); ROMK is known to be pH sensitive, exhibiting maximal channel opening at pH 7.8, and the majority of the channels are closed upon a shift to pH ∼6.6 (78, 79). However, the actual role of R311 in channel function remains disputed. pH gating was initially thought to be mediated by the formation of an RXR triad (R41, K80, R311) (69), but subsequent structural studies cast doubt on the formation of the triad, and instead favored a model in which R311 formed intermolecular salt bridges with E302 from an adjacent monomer (67). Finally, even though N377K appeared to exhibit wild-type-like growth, the yeast ODs vary greatly across the replicates (**Fig 3C**, error bar of dark purple line), which is consistent with stochastic toxic effects of this mutation (see below and **Discussion**). Ultimately, given their strong associations with Bartter syndrome phenotypes, we selected all five mutations to characterize further.

### A subset of disease-associated ROMK mutations destabilize the protein

Prior work established that potassium channel variants can lead to disease by interfering with channel conductance, open probability, or abundance at the cell surface (80). The last of these possibilities is regulated by cellular protein quality control pathways, which monitor the folding state of a protein both in the ER—which may lead to ERAD—or in later steps of the secretory pathway, which may lead to lysosome targeting (81–83). Since some of the identified mutations reside in the cytoplasmic domain (namely, G228E, T300R, R311W, L320P, and N377K), which includes the critical immunoglobulin-like region [see above and (15, 51)], we surmised that these mutations would decrease protein stability. To test this hypothesis, we measured protein stability via a cycloheximide chase analysis in *trk1*Δ*trk2*Δ yeast expressing each variant (84).

As shown in **Fig. 4A**, the G228E protein was highly unstable compared to wild-type ROMK, with almost no protein remaining after 60 minutes. These data are unsurprising given: (1) the severe growth defect observed in these cells after incubation in low potassium (**Fig 2**), (2) the change from glycine to a bulky charged amino acid (glutamic acid), which given the residue’s location in the β-sheet rich region might have drastic structural consequences, and (3) the Rhapsody score (0.930). Another mutation from TOPMed, L320P, which as indicated above resides in a β sheet at the base of this domain, also significantly destabilized the protein, almost to the same extent as G228E (**Fig 4A**; but see below). These results are consistent with its severe-to-moderate growth defects in yeast, the conversion of large hydrophobic residue to a structure-stabilizing proline, and a Rhapsody score that again predicts a deleterious outcome (**Fig 2B**, **S2 Fig** and **S3 Table**). It is also intriguing that the analogous residue (L321) in Kir2.1 resides in an amino acid patch (SYLANEI**<u>L</u>**W) that binds AP-1 and promotes Golgi export (85, 86), but the Golgi export consensus sequence is lacking in ROMK, suggesting instead that the structural change triggers premature degradation. In contrast to other variants, neither T86A nor T300R destabilized ROMK. In fact, the T86A and T300R substitutions appeared to stabilize ROMK (**Fig 4A** and see **Discussion**). For T86A, these results are consistent with its assignment as being neutral in Rhapsody (score: 0.063), and with robust growth observed in the yeast assay (**S1-S2 Figs** and **S3 Table**). On the other hand, it was somewhat surprising that the T300R protein was stable, despite a Rhapsody score of 0.722 and the moderate growth defect in yeast (**S1-S2 Figs** and **S3 Table**). We hypothesized that the mutation might instead alter a key architectural feature associated with ROMK function rather than overall stability (see below).

**Figure 4.**
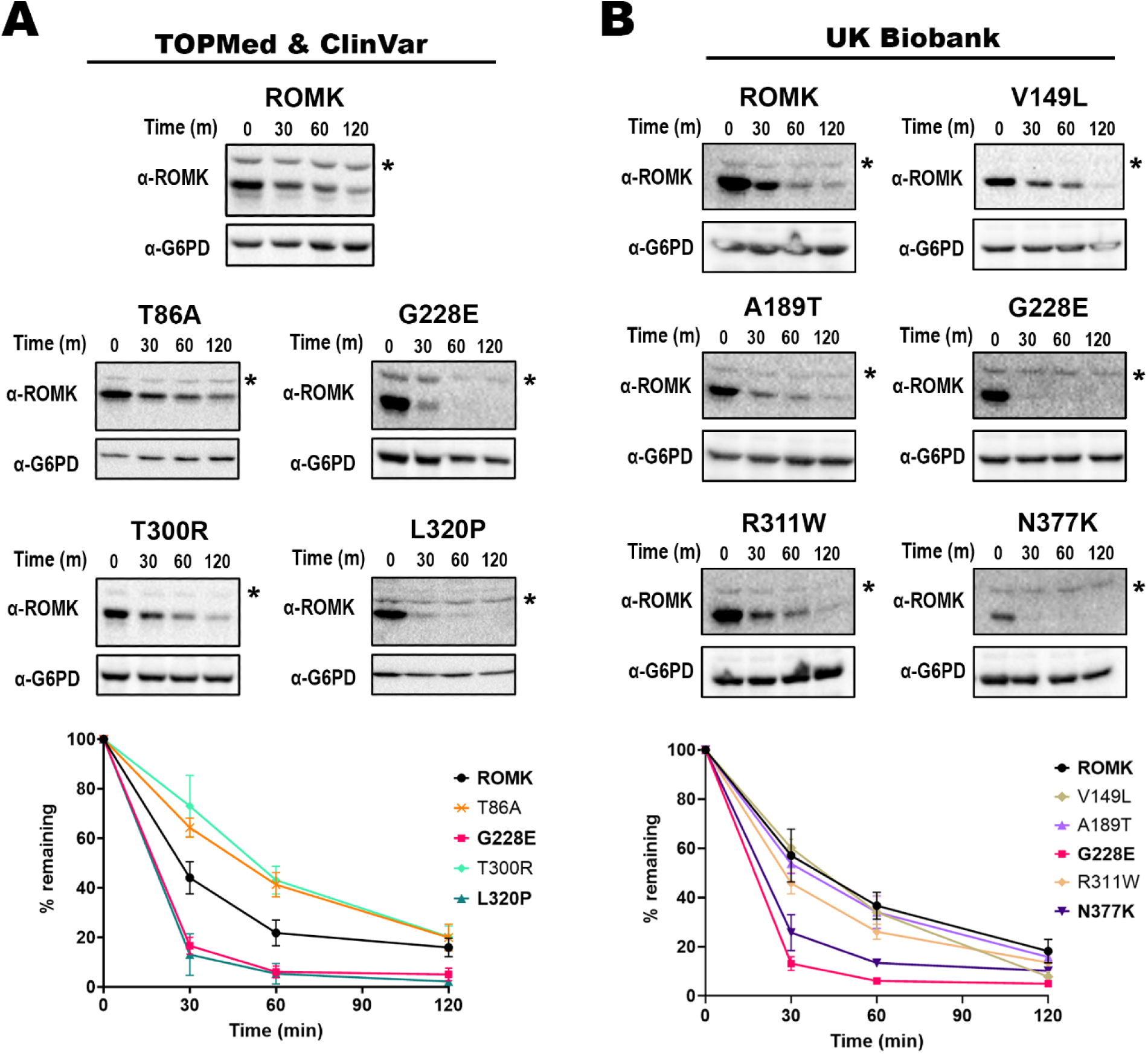
Select disease-linked mutations in the gene encoding ROMK destabilize the protein. Stability assays were performed in *trk1*Δ*trk2*Δ yeast expressing different ROMK variants from (A) TOPMed/ ClinVar and (B) the UK Biobank. In brief, yeast cultures were grown to mid-log phase, diluted, and incubated at 37°C for 30 min before cycloheximide was added. Cultures were then collected at the indicated time points and processed for immunoblot analysis. A rabbit antisera was used to detect ROMK (145), and a rabbit monoclonal antibody against G6PD was used as a loading control (see **Materials and Methods**). Representative immunoblots are shown, and graphs show the percentage of the protein remaining over time, compared to the 0 min (m) time point, as quantified using ImageJ (ver. 1.53c). * indicates a non-specific protein band recognized by the ROMK antisera. Graphs were made using GraphPad Prism (ver. 9.5.0), and data represent the means of at least three independent experiments, ± S.E. (error bars). For each experiment, a representative immunoblot is shown.

Among the UK Biobank mutations, only N377K—besides G228E—disrupted protein stability (**Fig 4B**), despite its lack of effect on yeast growth. Even though N377K did not result in a marked defect in yeast growth, there was a significant difference in growth phenotypes across replicates (**Fig 3C**), suggesting the acquisition of spontaneous suppressors (87) (also see **Discussion**). In any event, the net growth phenotype of N377K is consistent with its designation of a neutral mutation by Rhapsody (0.287).

We conclude first that the predictive power Rhapsody has been largely validated, with some exceptions. Specifically, 10 out of the 16 predicted deleterious mutations from both approaches exhibited growth defects in yeast, while all five predicted neutral mutations (N377K included) were without or with only minimal defects. Second, the accuracy of Rhapsody is enhanced for mutations with high Rhapsody scores, as evidenced by the fact that 8 out of 10 mutations with Rhapsody scores >0.7 impaired yeast growth. On the other hand, the correlation between Rhapsody scores and protein stability is weaker, which is perhaps unsurprising because a mutation can affect the structure at residues that alter channel conductance. Third, our results indicate that datamining efforts, across three human genomic databases, can identify previously uncharacterized ROMK mutations that appear to destabilize the protein, an outcome that may contribute to disease presentation and positions these mutations—and certainly other newly uncovered mutations—as targets of therapies that may one day restore ROMK folding, as seen for other protein conformational diseases (88, 89).

### Select ROMK mutants are targeted for ERAD and limit ROMK levels at the cell surface

Some ROMK variants that significantly alter structure (e.g., Y314C; **Fig 2**) are targeted for ERAD, as shown previously (15). Therefore, we next asked if the ERAD pathway is also responsible for the accelerated degradation rates observed in **Fig 4** for the G228E, L320P, and N377K alleles. To this end, we again performed stability assays in yeast, as described above, but measured protein turnover in the presence or absence of MG-132, a drug that inhibits a catalytic site in the proteasome (90). To prevent drug efflux, these experiments were performed in the *pdr5*Δ yeast strain that lacks a multidrug efflux pump (91), as extensively employed in previous studies [see e.g., (15, 50, 92)]. Consistent with ERAD targeting, protein stability assays revealed that all three mutant proteins were subjected to proteasome-dependent degradation (**Fig 5**, compare DMSO and MG-132 results). As shown previously (15), the wild-type channel was also targeted for proteasome-dependent degradation, but to a lesser extent (compare relative stabilities of wild-type versus the mutants in the DMSO control). This is likely due to the inefficient and inherently error-prone process of protein folding and channel assembly in the ER, and has been seen frequently for other ion channels, such as ENaC (93), CFTR (94), hERG (95), and Kir2.1 (50).

**Figure 5.**
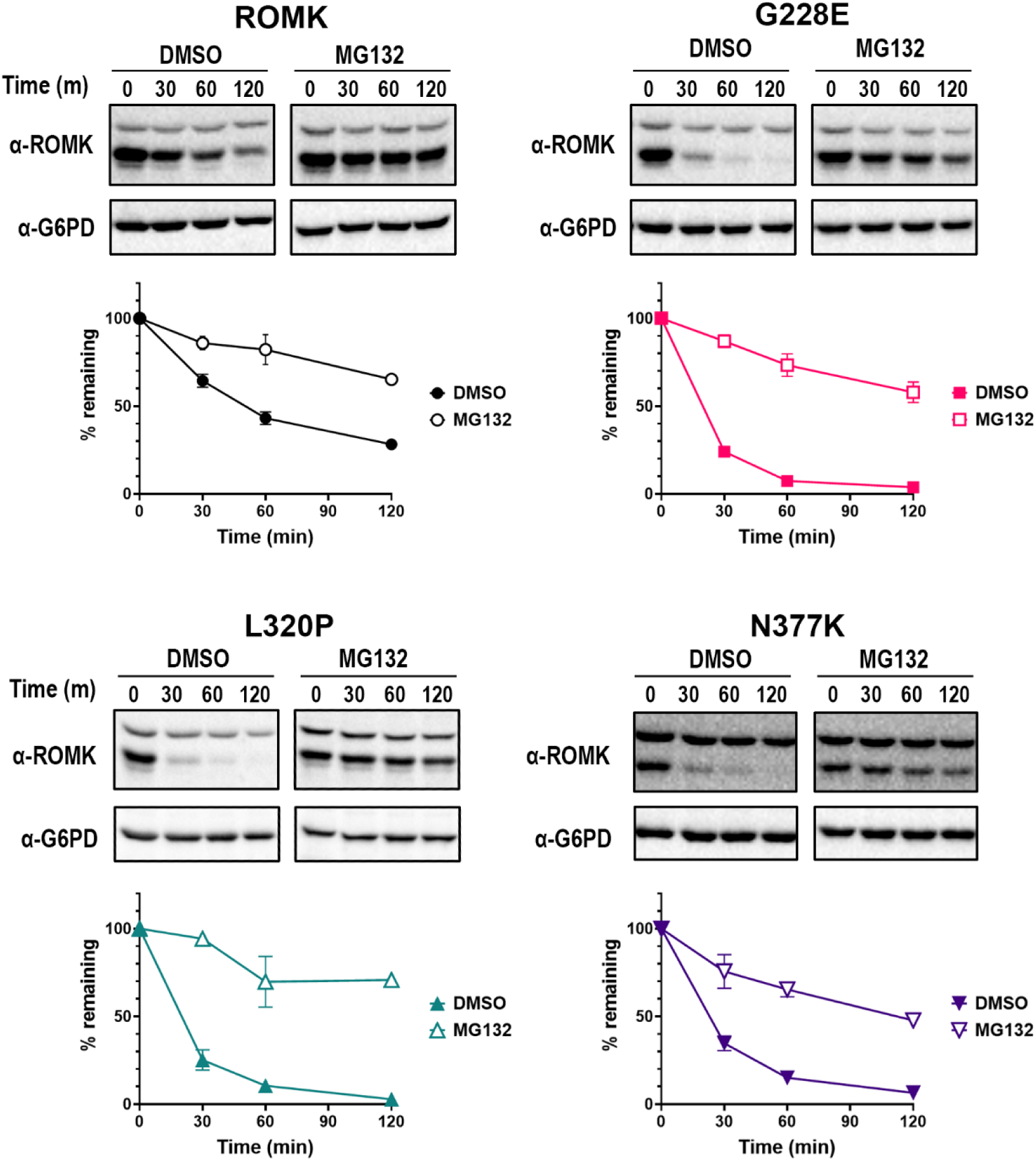
Three disease-associated ROMK mutants are degraded by the proteasome in yeast. Stability assays were performed in *pdr5*Δ yeast expressing wild-type ROMK, or ROMK carrying the G228E, L320P, or N377K mutation. Yeast cultures were grown to mid-log phase, diluted, and incubated at 37°C for 30 min with either the proteasome inhibitor MG-132 or an equal volume of the vehicle before cycloheximide was added. Cells were collected at the indicated times, processed, and immunoblot analysis was performed as described in the **Materials and Methods**. Representative immunoblots are shown, and graphs show the percentage of the protein remaining over time, compared to the 0 min (m) time point, as quantified by ImageJ (ver. 1.53c). Graphs were made using GraphPad Prism (ver. 9.5.0), and data represent the means of at least three independent experiments, ± S.E. (error bars). For each experiment, a representative immunoblot is shown.

To confirm that the three disease-associated mutants that underwent proteasome-dependent degradation are selected for ERAD, we next conducted stability assays in a temperature sensitive yeast strain, *cdc48-2* (96), which encodes a defective temperature-sensitive allele of *CDC48*, the gene encoding the AAA+-ATPase that mediates protein retrotranslocation in yeast (97, 98). At a non-permissive temperature, each of the ROMK mutant proteins was again significantly stabilized (**S4 Fig**).

To confirm our results in a more physiologically relevant cell system, we conducted stability assays in HEK293 cells transfected with each ROMK variant and again used MG-132 to inhibit the proteasome. In these assays, G228E, L320P, and N377K were degraded to variable extents when compared to wild-type ROMK (**Fig 6A**). Specifically, G228E and N377K exhibited somewhat higher degradation rates, with 55% and 37% of the protein remaining, respectively, by the end of the 4-hr experiment (compared to 59% for the wild-type protein). In contrast, L320P was only mildly unstable (again compare the relative curves in the presence of the DMSO control). In fact, the degradation rate of L320P (63% remaining) was nearly identical, if not slightly improved, compared to that observed for wild-type ROMK (see **Discussion**). Regardless, the degradation of each protein was slowed in the presence of MG-132.

**Figure 6.**
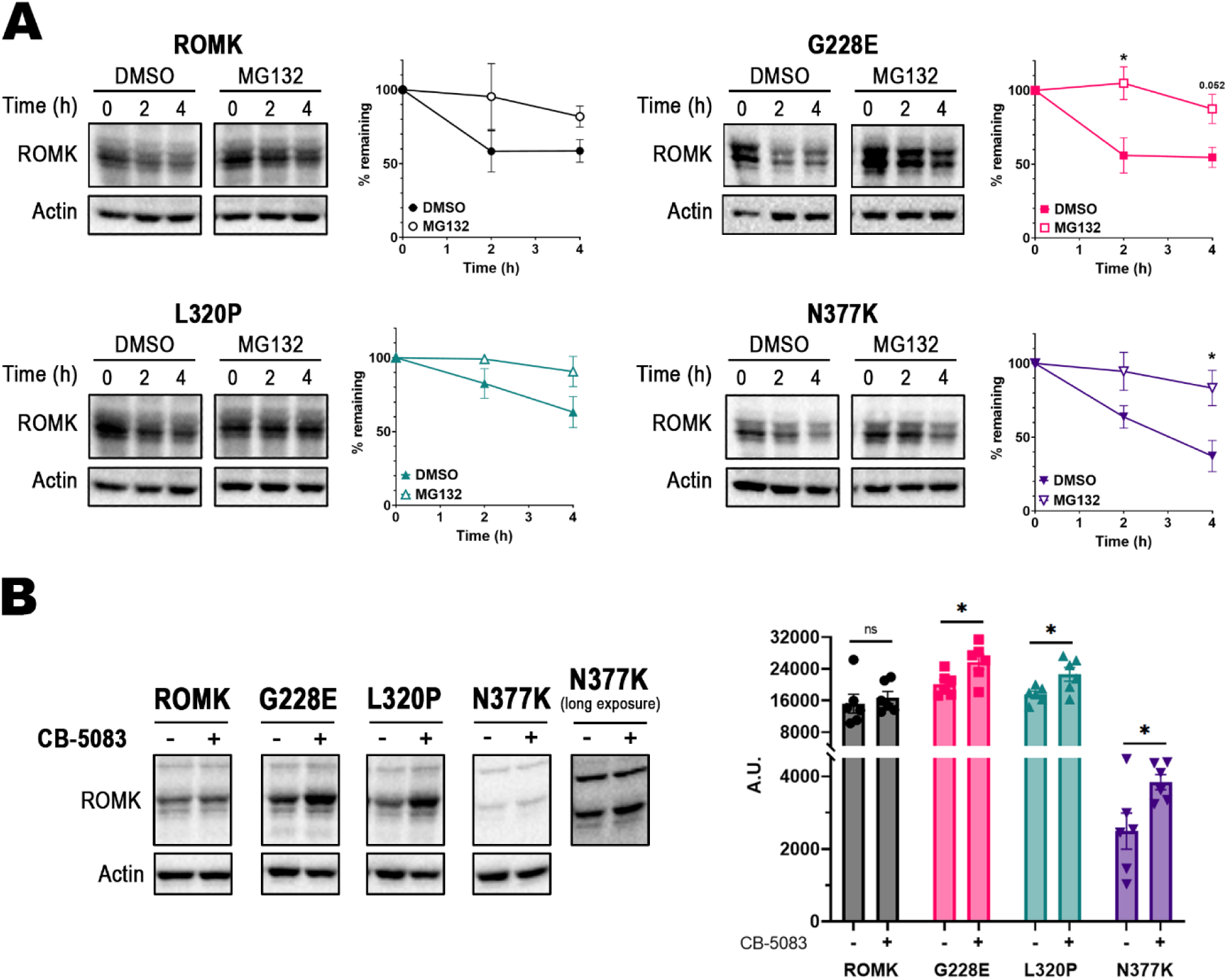
The ERAD pathway also contributes to ROMK mutant turnover in HEK293 cells. (A) Stability assays of HEK293 cells expressing wild-type ROMK, or ROMK carrying the G228E, L320P, or N377K mutation. HEK293 cells transfected with the indicated expression vector were treated with MG-132 or the equivalent volume of DMSO for 30 min, at which point cycloheximide was added. Cells were then processed as described in the **Materials and Methods**. Representative immunoblots are shown, and graphs show the percentage of the protein remaining over time, compared to the 0-hr (h) time point. A rabbit antisera was used to detect ROMK (145), and a mouse monoclonal antibody against actin was used as a loading control. (B) Steady-state protein levels before and after proteasome inhibition. The same cell lysis samples at timepoint 0 and 4 hr from (A) were reexamined by immunoblot analysis. (C) Steady-state protein levels before and after p97 inhibition with CB-5083 for 4 hr. Graphs were made using GraphPad Prism (ver. 9.5.0), and data represent the means of at least three independent experiments, ± S.E. (error bars). For each experiment, a representative immunoblot is shown, and the quantification was performed using ImageJ (ver. 1.53c); p-values in (B) and (C) were calculated with two-tailed Student’s t-test for independent samples. ns, p ≥ 0.05; *, p < 0.05.

We subsequently assessed ERAD in the presence or absence of an inhibitor of p97, which is the mammalian homolog of Cdc48 (99–101). The compound, CB-5083 (102), is somewhat toxic and has been used in clinical trials for various cancers (103). Thus, we performed steady-state measurements of ROMK after treatment with CB-5083 or the DMSO control, as employed previously (15). As shown in **Fig 6B**, a statistically significant increase in the G228E, L320P, and N377K mutant proteins was evident, whereas the levels of the wild-type protein in the presence or absence of CB-5083 were not as dramatically affected.

Although the N377K protein was unstable (**Fig 6A**), we routinely observed significantly less protein at steady-state and in the degradation assays at the 0 min time point (compare matched and long exposures of N377K and the other mutants, as well as the wild-type protein, in **Fig 6B**). Based on these results, we surmise that N377K ROMK either exhibits abortive translation or is rapidly degraded co-translationally, leaving a sub-pool that then turns over more slowly by ERAD. Each of these scenarios has been observed as a source of the molecular etiology underlying other diseases (104–106). While a full definition of this phenomenon awaits further analysis, this outcome might similarly give rise to Bartter syndrome type II.

It is important to highlight that the overall muted level of protein destabilization observed in HEK293 versus yeast cells is consistent with the fact that the ERAD pathway in yeast is hyperactive. Similar results with misfolded mutant alleles in ROMK and Kir2.1 have been observed previously (15, 50). It is also worth noting that the mutation with the most modest deleterious Rhapsody score (0.588), L320P, also exhibited the most wild-type-like degradation phenotype (**Fig 6A-B**; see **Discussion**).

The enhanced dependence on p97 to maintain the steady-state levels of the G228E, L320P, and N377K mutants in HEK293 cells suggests that lower levels of these proteins should reside at the cell surface. To test this hypothesis, we expressed the wild-type protein and the ROMK variants in HEK293 cells and performed cell-surface biotinylation assays to measure the plasma membrane protein pool (48, 107). As anticipated, markedly lower levels of biotinylated G228E and L320P channels were observed at the plasma membrane relative to the wild-type protein (**Fig 7**). In addition and as noted above, the levels of N377K were significantly lower in HEK293 cells, so the biotinylated protein pool at the cell surface was also drastically reduced (**S5 Fig**). As controls for labeling specificity, the Na+/K+-ATPase—a plasma membrane resident—was labeled and identified after avidin pull-down of the biotinylated material, whereas Hsp90, an abundant cytosolic protein, was absent. Taken together, these data indicate that disease-associated mutations identified from the complementary genomic databases deplete ROMK at the cell surface, which likely contributes to disease.

**Figure 7.**
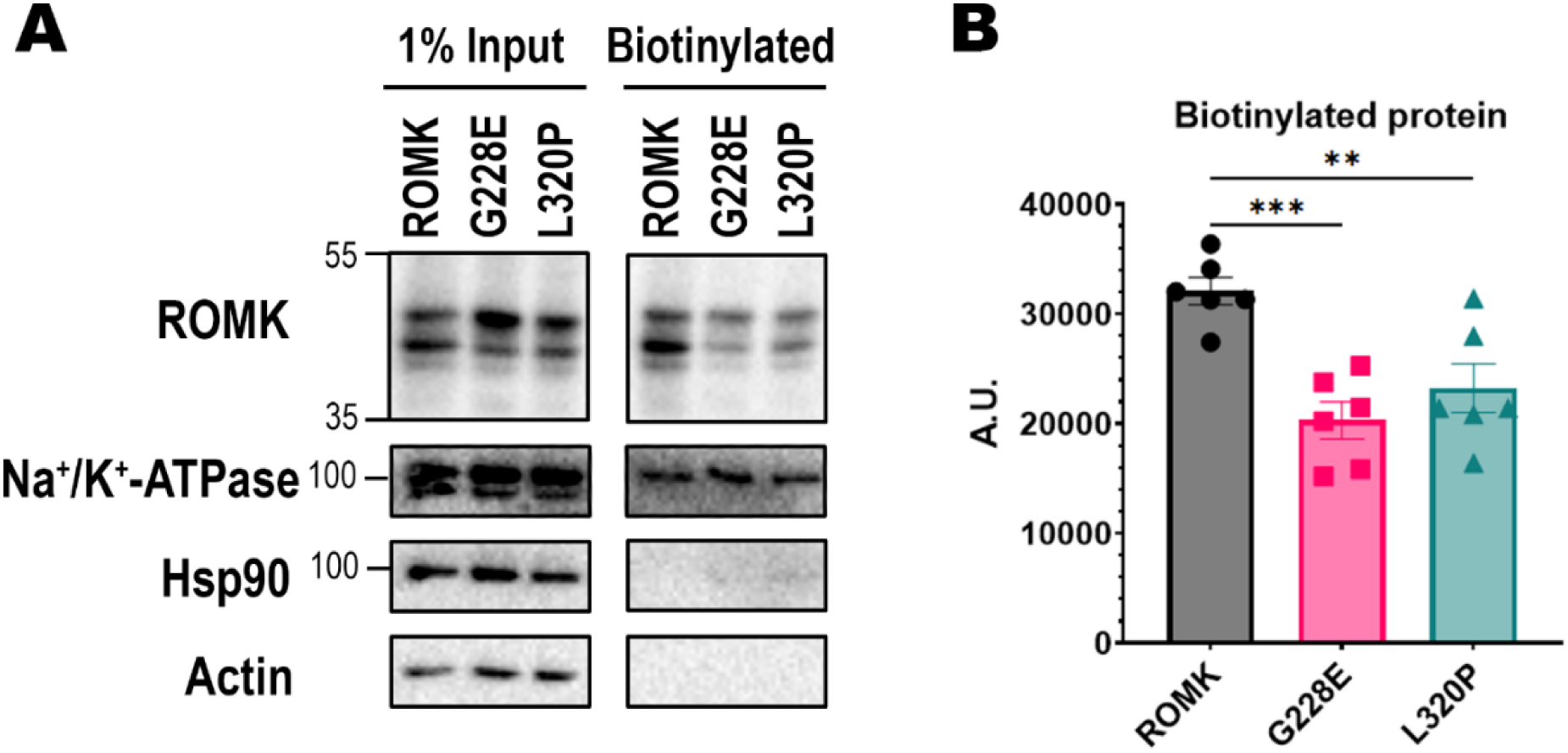
The cell surface levels of G228E and L320P ROMK are reduced in HEK293 cells. (A) A cell-surface biotinylation assay is shown of the relative surface expression levels of the indicated ROMK variants. HEK293 cells expressing wild-type ROMK or the G228E or L320P mutant were treated with biotin, processed, and incubated with streptavidin beads before an immunoblot analysis was performed. 1% input was collected prior to the overnight incubation, while the “Biotinylated” material represents precipitated protein. A representative immunoblot is shown, with a rabbit antisera to detect ROMK (145), a mouse monoclonal antibody against the Na+/K+-ATPase, a mouse monoclonal antibody for Hsp90, and a mouse monoclonal antibody against actin. (B) Graph shows the quantification for biotinylated protein, as measured by ImageJ (ver. 1.53c). Error bars represent the means of six independent experiments, ± S.E. p-values were calculated with two-tailed Student’s t-test for independent samples. ns, p ≥ 0.05; *, p < 0.05; **, p < 0.01; ***, p < 0.001.

### T300R abolishes channel activity

In contrast to changes in protein stability, protein deficiency, and/or altered abundance at the cell surface, disease-associated mutations in ion channels might traffic normally but are unable to support ion conductance, as observed for class III mutations in CFTR (108). Characterizing this phenotype is vital as—in contrast to the ERAD-targeted F508del CFTR protein repaired by chemical chaperones—the class III mutant defects can be treated with approved potentiators (109, 110). For ROMK, the mechanisms of channel gating are still being actively investigated (9, 41), but the general consensus is that channel gating capitalizes on the helix-bundle crossing region, PIP_2_ binding, and a narrow opening at the top of the cytoplasmic pore, known as the G-loop, as described above (9, 40). Therefore, we focused on one of the mutations identified from the ClinVar database, T300R, which is located on the G-loop. As shown above, this mutant compromised the growth of the *trk1*Δ*trk2*Δ yeast strain in low potassium (**S2 Fig** and **S3 Table**), suggesting defective potassium transport, but the protein was stable (**Fig 4**). Similar observations were made in a previous study in which two ROMK mutations, P185S and R188C, moderately increased protein surface expression, yet negatively affected channel gating and conductance in a PIP_2_-dependent manner (44). We thus predicted that T300R would reduce channel currents, as a homologous mutation in the closely related Kir2.1 channel, M301R, prevented channel function (42). In addition, modeling of T300R onto the structure suggests that the change from a small hydroxyl into a large basic side chain likely occludes the cytoplasmic pore and prevents potassium passage (**S6 Fig**).

To measure channel activity, we performed two-electrode voltage clamp assays in *Xenopus* oocytes expressing wild-type and select ROMK variants. As hypothesized, the T300R mutation completely abolished ROMK current **(Fig 8A-B)**, reducing ROMK- specific, i.e., barium sensitive, currents to the same level as the negative controls (water-injected as well as the Y314C mutant [see above and (14, 15, 51)]). Because another mutation at the same site, T300I, was one of the 17 alleles identified from the TOPMed and ClinVar databases (**Fig1** and **S1-S2 Tables**), we also examined currents corresponding to this variant. In contrast to the effect of the T300R allele, the current was identical to wild-type ROMK when oocytes were injected with a cRNA for T300I ROMK. The wild-type-like current is perhaps expected given the lack of a growth defect in T300I-expressing yeast (**S2 Fig** and **S3 Table**) as well as the less consequential substitution from one to another beta-branched amino acid. Thus, these results shed light on a functional defect associated with a stable Bartter syndrome-associated mutation in the gene encoding ROMK. Moreover, these data highlight the power of uniting a computational analysis and the yeast system as an initial read-out to screen ill-characterized and previously undefined alleles in a potassium channel encoding gene.

**Figure 8.**
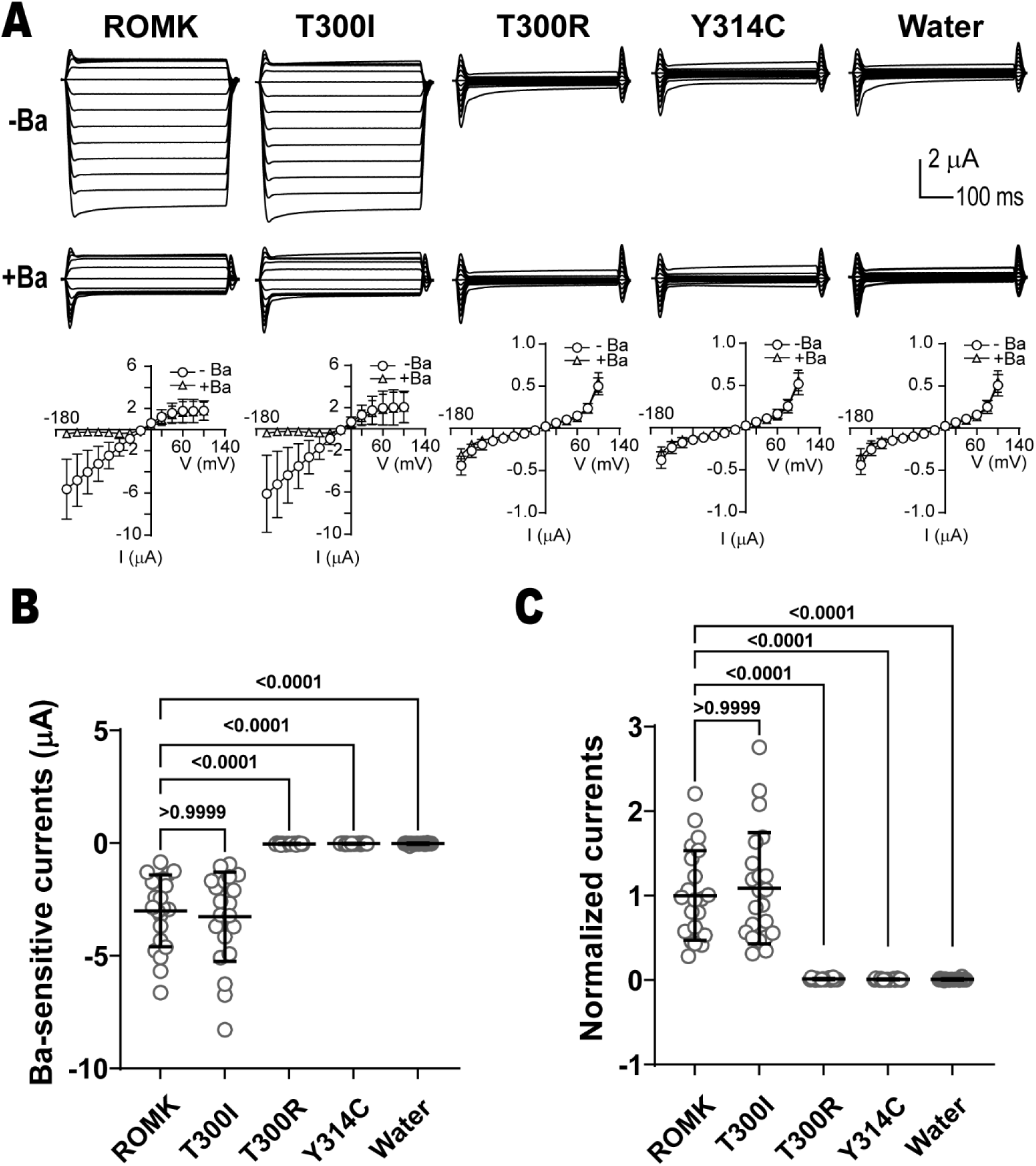
The T300R mutation in ROMK abolishes channel currents. (A) Top panel: Currents recorded by two-electrode voltage clamps (TEVC) in *Xenopus* oocytes. Oocytes from female *Xenopus laevis* were injected with 1 ng of the indicated cRNAs, or the equivalent volume of water. 20-30 hr following cRNA injection, TEVC recordings were measured at different voltages (-160 mV to 100 mV, in 20 mV increments) in a bath solution containing 50 mM KCl (for more details, see **Materials and Methods**). Currents were recorded in the presence or absence of 1 mM BaCl_2_ and I-V plots are shown (bottom). In addition to a water-injected control, the results with a known unstable disease-causing mutant (Y314C) are shown (15, 51). (B) Graph shows the Ba2+- sensitive/ROMK current in oocytes injected with the indicated conditions, as recorded by TEVC. (C) Normalized currents, which are defined as Ba2+-sensitive currents divided by the means of the wild-type currents. Error bars in the graphs in (B) and (C) show the means of 22 replicates, ±S.D. p-values (shown above the data) were computed using Kruskal-Wallis and Dunn’s multiple comparisons tests. The data shown are a representative result of three independent experiments using three batches of oocytes.

## Discussion

In the kidney, efficient plasma filtration and electrolyte reabsorption are achieved through a system of transporters and ion channels (111), among which ROMK plays a crucial role. Potassium efflux through ROMK in the thick ascending limb and the cortical collecting duct of the kidney nephron helps maintain potassium and sodium homeostasis (9, 112). Over 40 missense mutations in the gene encoding ROMK, *KCNJ1*, have been identified and linked to Bartter syndrome type II (4), a rare autosomal recessive disease presenting with fluid loss and electrolyte imbalance, i.e., renal salt wasting, polyuria, early post-natal hyperkalemia and subsequently hypokalemia (113). Previous investigations of the cellular mechanisms of Bartter-associated ROMK mutations have primarily focused on variants that affect whole-cell currents (12, 114), and each study commonly analyzed a handful of mutations. Thus, for both this disease and most other protein conformational diseases, there exists a need to establish a high-throughput method to systematically identify potential disease-causing variants in the genome, especially with the increasing availability of human genomic and phenotypic data from large-scale worldwide studies (115).

In this study, we utilized two computationally-guided approaches to mine three human genomic databases (TOPMed, ClinVar, and the UK Biobank), with the ultimate goal of identifying novel and previously uncharacterized mutations associated with Bartter syndrome type II. From the initial analyses, 21 mutations were selected for expression and functional screening in the established *trk1*Δ*trk2*Δ yeast system (24, 25), among which one mutation (G228E) was identified from both approaches. Based on results from yeast viability assays, we again validated the ability of the Rhapsody algorithm to assess mutation severity, as the growth of yeast carrying the mutant proteins reflected their Rhapsody scores 71% of the time (i.e., 15 out of 21 mutations exhibited growth phenotypes in accordance with Rhapsody scores: 10/16 deleterious and 5/5 neutral). As highlighted in the **Results**, Rhapsody was more effective for scores >0.7, with an accuracy of 80%, compared to 62.5% when all predicted deleterious mutations were considered. Therefore, future studies employing this method for mutation assessment should exercise caution for lower-scoring amino acid substitutions.

Because most prior functional analyses of ROMK variants focused on those that impair channel function (12, 114), we specifically sought mutations that compromise protein folding and trafficking. To this end, we conducted functional assays in yeast, *Xenopus* oocytes and HEK293 cells, thereby revealing distinct cellular mechanisms underlying potential disease etiology. One newly identified and previously uncharacterized Bartter mutation (T300R) had no effect on protein stability but blocked channel conductance. In contrast, three mutants (G228E, L320P, and N377K) were highly unstable in yeast, but exhibited varying degrees of stability in mammalian cells. G228E, which was uncovered from both screening strategies, likely affected protein folding due to the substitution of a small, aliphatic amino acid to a larger charged residue. This effect is also consistent with the mutation’s localization in the β sheet-rich cytoplasmic domain (see above). Despite decreased difference in protein degradation rates between the wild-type protein and the G228E mutant in mammalian cells, cell surface expression was significantly affected. Perhaps unsurprisingly, this is consistent with our finding that two individuals heterozygous for G228E from the UK Biobank exhibited issues with their amniotic cavity and membrane, a typical manifestation of antenatal Bartter syndrome (70, 113). It is also important to note that more pronounced protein destabilization in yeast likely results from the hyperactive ERAD pathway in this organism, as described previously when ROMK and Kir2.1 were examined (15, 50).

Another Bartter mutation of uncertain clinical significance, L320P, also destabilized the protein in yeast, yet there was little effect on stability in HEK293 cells. Since this mutation is also located in the immunoglobulin domain, where thus far seven ROMK mutations can compromise protein stability, we reasoned that the folding of this domain in the ER is a rate-limiting step, at least in yeast. Despite the lack of an effect on protein stability in mammalian cells, we still observed a significantly reduced protein pool on the cell surface. This fact, coupled with its wild-type-like protein level at steady-state, suggest that the L320P mutation compromises ROMK trafficking at later steps in the secretory pathway, a process that may be rate-limiting in higher cells (116).

While the third mutation, N377K, initially appeared to lack a growth defect in yeast, there were significant stochastic effects between experiments (note the larger error of these measurements in **Fig 3** compared to the other strains). Nevertheless, the mutant protein was rapidly degraded in yeast through the ERAD pathway, and to a lesser extent in mammalian cells. Curiously, Rhapsody designated a neutral score for this mutation (0.287), perhaps reflecting the limitation of this program in analyzing mutations that rely on structural predictions (i.e., AlphaFold) instead of homology models. It is also possible that the amino acid substitution alters a critical post-translational modification or allostery, which cannot be captured by Rhapsody. This possibility is supported by the discovery of a nearby residue, N375, that resides within a non-canonical endocytic signal (Yx<u>N</u>PxFV) that binds to the ARH adaptor and recruits ROMK to clathrin-coated pits (117). This hypothesis is consistent with our proposal, above, that later steps in the trafficking pathway are altered.

Although with some faults, the *trk1*Δ*trk2*Δ yeast model provides a rapid, inexpensive, and quantitative route to screen mutations that affect potassium channel folding, trafficking to the cell surface, and function. Because this system is also amenable for drug discovery (25), future work will attempt to rescue variants whose defects were confirmed in higher cells (e.g., G228E). Yet, discrepancies between defects in yeast growth, protein stability, and/or confounding results in higher cells—as seen for N377K—hint at variables that must be taken into account in future screens. Based on the growth of transformants on plates that displayed a range of colony sizes, as well as the larger errors seen in growth assays (see above), the N377K mutation may cause toxic effects on yeast growth, which results in the accumulation of spontaneous suppressors (87) and the formation of “petite” colonies (see for example (118, 119)). Indeed, spontaneous suppressors arising from mutations in hexose or amino acid transporters are observed as common causes for phenotypic reversion in *trk1*Δ*trk2*Δ yeast (120–122). To test the latter possibility, we propagated cells from colonies of yeast expressing the N377K mutant on medium containing a nonfermentable carbon source (i.e., glycerol) instead of glucose. We found that the smaller colonies failed to grow on plates containing glycerol (**S7 Fig**), a phenotype typical of petite yeast (123) that occurs due to spontaneous mutations in or the loss of the mitochondrial DNA (124, 125).

To mine the UK Biobank data, we utilized REVEAL: Biobank, a high-performance, cost-effective computational platform for exploring, querying, and analyzing multi-omic biobank-scale datasets (35–38). REVEAL: Biobank’s ability to rapidly filter a large search space to create cohorts of interest, execute complicated bioinformatics workflows at scale in a user-friendly manner, and allow custom algorithms (e.g., phecode generation) to be easily applied positions REVEAL: Biobank as an optimal solution for high-throughput *ad hoc* analysis. Moreover, multiple algorithms, such as SAIGE (64) and REGENIE (65), can be incorporated into the workflow with simple parameter changes. This allows results to be validated, which is vital given discrepancies frequently observed in bioinformatics tools.

In addition to the correlative associations obtained from GWAS, the beta value, i.e., effect size, provides a powerful measure of the degree and direction of impact that a mutation has on the phenotype. While more work is needed to verify the degree of the impact observed, the direction of the beta values (+/-) in **Table 1** follows the direction of the difference seen in the mean values of the phenotypes between the wild-type and mutant genotype cohorts in **Table 3**.

Results obtained from the UK Biobank GWAS analyses largely corroborated the findings from the yeast screen and Rhapsody predictions, but it is interesting to note that the p-value of the associations were lower than those deemed significant in typical GWAS (1e-08). This highlights the need to have multiple approaches of hypothesis testing and validation, along with potential limitations of *in silico* models. Future efforts might also utilize metrics other than the p-value to determine significance (126). A probable explanation for the low p-values could be the high imbalance observed in the ratios of cases (individuals with a phenotype) and controls, and of wild-type and mutant genotypes (see **Table 3** and **S5 Table**).

The pipeline using REVEAL: Biobank described in this paper can also be expanded into two directions to further dissect the cellular and biochemical mechanism underlying ROMK function and to elucidate the relationship between ROMK and other diseases. In the first direction, we can use other known phenotypes associated with ROMK, such as hypertension, to uncover additional mutations that exert a functional effect on trafficking or function. This approach can also be extended to include linkage disequilibrium calculations coupled with burden tests to identify co-occurring mutations in proteins known or thought to interact with ROMK, essentially identifying synthetic interactions, but not necessarily synthetic lethal interactions (127, 128). We previously obtained these outcomes with the cytoskeletal scaffold protein encoded by *SLC9A3R2* (129). Finally, the entire set of mutations could be fed into an artificial intelligence (AI)-based application, such as the AlphaFold Protein Structure Database (75), to provide insights into the structural implications of the mutations. The second direction is a bootstrapping approach to uncover potential new disease connections by leveraging both genetic and health record data to explore longitudinal prescription and general practitioner information (i.e., READ codes) for patients with identified mutations in key genes (130–132). Consequently, there is ample opportunity to further explore ROMK/*KCNJ1*, and other disease-linked genes, by leveraging large human datasets in the UK Biobank.

In sum, our work highlights a pipeline for computational-guided mining of human databases to search for mutations in any potassium channel that can be assayed in the yeast model. We identified and uncovered the cellular mechanisms underlying disease phenotypes in a subset of ROMK mutations with uncertain clinical significances, among which three destabilize the protein, while one is channel-defective. It is worth noting, however, that uncharacterized ROMK mutants remain, and new disease-associated variants will continue to arise. Future work should thus focus on improving the high throughput nature and signal-to-noise of the yeast assay so that more mutations can be simultaneously screened, which combined with studies in higher cells may ultimately contribute to the development of precision medicine to treat those with Bartter syndrome type II.

## Materials and Methods

### Computational analysis and selection of mutation from the TOPMed & ClinVar databases

At the time of this study, data from the Trans-Omics for Precision Medicine (TOPMed) program (28) were publicly available in its “freeze 5” version on the Bravo server (39). This version of the dataset consists of 463 million variants from 62,784 individuals and that specifically contains 758 genomic variants and 124 predicted missense mutations in *KCNJ1*. To analyze the potential severity of the mutations, we ran a saturation mutagenesis analysis of ROMK with Rhapsody (30, 31) available on a web interface (http://rhapsody.csb.pitt.edu/). We used a homology model of human ROMK (Uniprot #: P48048) obtained from Swiss-Model (133), which was built based on the crystal structure of Kir2.2 (PDB ID: 3SPG) (134). Thus, Rhapsody was able to compute the pathogenicity probability, i.e., “score”, only for amino acid residues 38-364 that are available in the homology model. Because N377 is absent from this model, a Rhapsody score for the N377K mutation was obtained using a monomeric structure predicted by AlphaFold (75). For further analysis, we prioritized mutations with a high Rhapsody score, i.e., more deleterious, as well as mutations located in regions previously found to be important for protein folding and channel function (see text for additional details). We also focused on mutations associated with Bartter syndrome that were classified clinically as being “of uncertain significance” in the ClinVar database (29). Thus, T300R from ClinVar was added based on this classification, and also due to its position at residue T300 (since T300I had been selected from TOPMed).

### GWAS analysis on data from the UK Biobank

As noted in the **Results**, whole exome sequencing (WES) data from the UK Biobank (32) was used to perform three genome-wide association studies (GWAS). At the time of this analysis, the WES data had been released for ∼200,000 individuals out of the ∼500,000 total UKBB participants (56, 57), among which there are 511 *KCNJ1* variants (**S4 Table**). After applying a minor allele frequency (maf) cutoff of >1e-5, the number of mutations used for the GWAS was 142. Phenotypic data were selected from the pool of the ∼200,000 participants and included: (1) 15 continuous/ quantitative phenotypes for relevance to ROMK function, Bartter syndrome type II, and hypertension (9, 63) (e.g., systolic and diastolic blood pressure, serum urea, creatinine, calcium, and phosphate, and urine potassium and sodium), (2) 15 phenotypic codes, or “Phecodes” (34), associated with Bartter syndrome type II, and (3) 168 continuous/quantitative metabolomics biomarkers . The quantitative phenotypes were normalized using inverse rank transformation to address non-normality of the underlying distribution (135).

The phecodes that were chosen represented 25 unique ICD10 codes relevant to Bartter syndrome, but individuals with phecodes related to diabetes and hypothyroidism were excluded from the analysis (see **Results**). Phecodes can be described as a mapping of grouping International Classification of Diseases (ICD) codes into clinically relevant groups (34, 136). Phecodes improve the power for association studies and enhance the accuracy of relevant phenotypes, in contrast to ICD codes. Specifically, we developed a custom algorithm to generate phecodes relevant to Bartter Syndrome based on an unsupervised multimodal automated phenotyping method (137). The metabolomics biomarkers from the UK Biobank (data field category 220) were measured in plasma samples using a high-throughput NMR-based metabolic biomarker profiling platform developed by Nightingale Health Ltd.

The GWAS analyses were done using REVEAL: Biobank, a computational platform designed to explore, query, and perform large computations on biobank-scale datasets (35–38). REVEAL: Biobank comprises R and Python application programming interfaces (API) for programmatic access to data and graphical user interfaces (GUI’s) for selection of cohorts using phenotype and genotype filters, and then analyzes GWAS and Phenome Wide Association Studies (PheWAS) results from a browser window. REVEAL: Biobank is built upon SciDB (51), a database solution ideal for storing and querying multi-omics data, utilizes elastic scaling through an application called BurstMode for efficient and cost-effective analyses and flexFS, a networked POSIX compliant filesystem for working with big data. REVEAL allows rapid and FAIR (a group of guiding principles for scientific data management (138)) access to the UK Biobank data, and multiple users can load, read, and write data in a secure, transactionally safe manner as data operations are guaranteed to be atomic and consistent (ACID compliant).

We used two algorithms, SAIGE (v0.44.6.5) (64) and REGENIE (v2.0.2) (65), to carry out the association analyses. Both algorithms are standards in GWAS bioinformatics workflows and are used to perform a regression test between a mutation of interest and a phenotype. Utilizing two algorithms also helped validate results. There were 12 covariates used in the GWAS: age, sex, and 10 genetic principal components provided by the UK Biobank (data field 22009).

The selection of alleles for further characterization is described in the **Results**.

### Plasmid construction

Rat ROMK1 was amplified from the pSPORT1-ROMK1 vector (139) and inserted into the yeast expression vector pRS415 with SmaI and XhoI and was flanked by the *TEF1* promoter and *CYC1* terminator (140), as described (15, 31). Point mutations in *KCNJ1* were introduced into the resulting pRS415TEF1-ROMK1 vector using either two-step overlap extension mutagenesis (141) or site-directed mutagenesis with the QuikChange kit (Agilent Technologies, CA, USA, catalog # 200523). To express ROMK variants in HEK293 cells, the DNA inserts were digested with BamHI and XhoI from the yeast vector and subcloned into pcDNA3.1(+). The DNA sequences of all variants in the ROMK inserts were confirmed by Sanger sequencing (GENEWIZ, S Plainfield, NJ, USA). All primers used in this study are listed in **S6 Table**.

### Yeast strains and growth conditions

A *Saccharomyces cerevisiae* strain lacking the Trk1 and Trk2 potassium transporters, *trk1*Δ*trk2*Δ, was employed to assess mutation severity by measuring the ability of each mutation to restore growth on low potassium medium, as described previously (15, 31, 48, 50). Briefly, plasmids were transformed into yeast via the standard lithium-acetate method (142), and yeast were grown at 30°C in liquid or solid synthetic complete (SC) medium lacking leucine, which contained monosodium glutamate as the main nitrogen source and buffered to pH 4.5 with MES. Media was supplemented with either 100 mM or 25 mM KCl. Due to the presence of residual potassium in the agar and nitrogen source, each plate contained an additional 7-10 mM KCl (46, 143).

To perform protein stability assays in yeast (see below), we utilized the indicated yeast strains (i.e., *trk1*Δ*trk2*Δ and *pdr5*Δ; see **S6 Table**). Cells were grown at 30°C and switched to 37°C at the beginning of the chase. Assays using *CDC48* and the isogenic *cdc48-2* strains were propagated at 26°C and then shifted to 39°C (**S6 Table**).

### Yeast viability assays

Yeast viability assays were conducted as described (31). For serial-dilution growth assays on solid medium, saturated overnight cultures were diluted to an A_600_ of 0.20, then further diluted 5-fold four times in a standard 96-well plate, followed by inoculation into SC-Leu medium supplemented with 100 mM or 25 mM KCl using a 48-pin replica plater (Sigma-Aldrich, St. Louis, MO, USA). Plates were incubated at 30°C and imaged after two days with the Bio-Rad ChemiDoc XRS+ imager. For assays in liquid medium, saturated overnight cultures were diluted to an A_600_ of 0.20 with SC-Leu medium containing 25 mM KCl in a 96-well plate. The plates were then covered with a Breathe-Easy gas permeable membrane (Diversified Biotech, Dedham, MA, USA), and cell density readings were recorded using the Cytation 5 plate reader (BioTek, Winooski, VT, USA) every 30 min for the indicated time with constant shaking at 30°C.

### Yeast stability assays

Stability assays in yeast were carried out based on established protocols (15, 50), with minor modifications. In brief, yeast cultures transformed with the ROMK expression vector (see above) were grown in selective media to mid-log phase (A_600_ = 0.7 – 1.5), diluted to the same density (typically A_600_ = 1.0), and transferred to a water bath with constant shaking at 200 rpm. The cells were then incubated for 30 min at 30°C (*trk1*Δ*trk2*Δ) or for 2 hr at 39°C (the *CDC48* and *cdc48-2* strains). A similar protocol was followed for when the *pdr5*Δ strain was employed, except the initial 30 min incubation was performed in the presence of 50 µM MG-132 or an equal volume of the vehicle (DMSO).

Next, cycloheximide was added to a final concentration of 150 µg/ml, at which point a 1 ml aliquot was collected. Subsequent 1 ml aliquots were collected at the indicated time points, flash frozen in liquid N_2_, and either kept at -20°C or were subject to immediate processing and lysis.

The levels of ROMK at each time point were assayed as previously outlined (15, 48). After lysis in 300 mM NaOH, 1% β-mercaptoethanol, 1 mM PMSF, 1 µg/ml leupeptin, and 0.5 µg/ml pepstatin A, total protein was precipitated with 5% trichloroacetic acid on ice. The mixture was the centrifuged at 14,000 rpm for 10 min at 4°C in a microfuge and subject to SDS-PAGE and immunoblot analysis. See **S6 Table** for more information on the antibodies and dilutions used.

### HEK293 cell culture, transfection, and stability assays

HEK293 cells (Thermo Fisher, Waltham, MA, USA) were cultured at 37°C in Dulbecco’s Modified Eagle’s Medium containing high levels of glucose (Sigma-Aldrich, St. Louis, MO, USA) and supplemented with 10% Fetal Bovine Serum and a mixture of penicillin/streptomycin (final concentration: 500 units/ml). Cells in 6-well dishes (passage 2-3, 60-90% confluency) were transfected with 2 µg of plasmids carrying the indicated ROMK mutants using Lipofectamine 2000 (Invitrogen, Waltham, MA, USA), and the media was replaced after 4 hr. Protein stability was measured based on an established protocol, with slight modifications (15). In short, fresh media containing 50 µM MG-132 or the equivalent volume of DMSO was added 18-20 hr post transfection. After a 30 min incubation, a final concentration of 50 µg/ml cycloheximide was introduced, and cells were collected at the indicated time points. For steady state measurements after treatment with CB-5083, a slightly modified protocol was followed. In brief, HEK293 cells were cultured in 12-well dishes and transfected with 0.6 µg of the indicated ROMK expression vector, and 20 hr post transfection, the media was replaced with media containing 50 µg/ml cycloheximide in the presence or absence of a final concentration of 20 µM CB-5083. Cells were collected after a 4 hr incubation at 37°C. In both assays, cell pellets were collected and retained at -20°C.

Cells were lysed in TNT buffer (50 mM Tris, pH 7.4, 150 mM NaCl, 1% Triton X-100) supplemented with a protease inhibitor cocktail (Roche, Basel, Switzerland) on ice for 20 min with occasional agitation. The mixture was then centrifuged at 13,000 rpm for 10 min at 4°C in a microfuge to remove the nuclear fraction, and the supernatant was transferred into new tubes and SDS sample buffer containing 150 mM DTT was then added to facilitate protein analysis by SDS-PAGE and immunoblots, as described (15, 48).

### Cell-surface biotinylation assays

Cell-surface biotinylation assays were performed as published (48), with minor modifications. In short, 20-22 hr post transfection, HEK293 cells expressing the indicated ROMK construct were treated with a final concentration of 125 µg/ml cycloheximide for 2 hr at 37°C. The plates were then transferred onto ice, washed three times, and treated with 0.3 mg/ml EZ-Link Sulfo-NHS Biotin (Thermo Fisher, Waltham, MA, USA) for 1 hr. Excess biotin was quenched by washing the cells with 100 mM glycine two times, and then the cells were lysed in 20 mM HEPES, pH 7.6, 1 mM EDTA, 1 mM EGTA, 25 mM NaCl, 1% Triton-X, 10% glycerol containing a protease inhibitor cocktail (Roche, Basel, Switzerland) for 1 hr before the mixture was centrifuged cold at 14000 rpm for 15 min to remove any insoluble material. The concentration of the liberated soluble protein was assessed with the Pierce BCA protein assay kit (Thermo Fisher, Waltham, MA, USA), and equal amounts of protein (180-250 µg) were brought to a total volume of 1 ml in the same buffer as above. After an aliquot corresponding to 1% of the total was collected, the remaining protein was added to 30 μl of Pierce NeutrAvidin-agarose beads (Thermo Fisher, Waltham, MA, USA) and incubated overnight at 4°C. The next day, the beads were washed three times and subject to SDS-PAGE and immunoblot analysis (see **S6 Table**).

### Two-electrode voltage clamp measurements

pRS415-ROMK expression plasmids (see above) were linearized and used as templates for cRNA synthesis by *in vitro* transcription using T7 RNA Polymerase (Ambion, Inc., Life Technologies, Carlsbad, CA, USA). The resulting cRNAs were then purified with an RNA purification kit (Qiagen, Hilden, Germany), quantified, and the cRNA quality was assessed by denaturing agarose gel analysis.

Oocytes from *Xenopus laevis* were harvested with a protocol approved by the University of Pittsburgh’s Institutional Animal Care and Use Committee. Briefly, stage V and VI oocytes were treated with collagenase type II and trypsin inhibitor to remove the follicle cell layer. Oocytes were then injected with 1 ng of the indicated cRNA and incubated at 18°C in a slightly modified Barth’s solution (15 mM HEPES, pH 7.4, 88 mM NaCl, 10 mM KCl, 2.4 mM NaHCO_3_, 0.3 mM Ca(NO3)_2_, 0.41 mM CaCl_2_, 0.82 mM MgSO_4_, 10 µg/ml streptomycin sulfate, 100 µg/ml gentamycin sulfate) for 20-30 hr. Next, two electrode voltage clamp experiments were performed at room temperature (20-24°C) with the TEV200A Voltage Clamp Amplifier (Dagan Corporation, Minneapolis, MN, USA) and the DigiData 1440A and Clampex 10.4 software (Molecular Devices, San Jose, CA, USA). Oocytes were placed in a recording chamber and perfused with a bath solution (10 mM HEPES, pH 7.8, 50 mM KCl, 48 mM NaCl, 2 mM CaCl_2_, 1mM MgCl_2_) at a constant flow rate of 5-10 ml/min. Whole-cell currents were recorded at a series of voltages (-160 and 100 mV in 20 mV increments), in the absence and presence of 1mM BaCl_2_ in the bath solution. Data were analyzed using Clampfit in the pClamp 10.4 package. Ba2+-sensitive currents, which represent ROMK channel activity in oocytes (144), were defined as the difference in currents measured in the absence and presence of BaCl_2_. Ba-sensitive currents were quantified, and graphs were made using GraphPad Prism (ver. 9.5.0).

### Statistical Methods

For the stability assays, CB-5083 treatments and cell-surface biotinylation in HEK293 cells, p-values were calculated with a two-tailed Student’s t-test for independent samples. In the two-electrode voltage clamp experiments, statistical analysis was conducted using Kruskal-Wallis and Dunn’s multiple comparisons tests, and normality was examined with Shapiro-Wilk tests.

## Supporting information

All supplemental information (except S4 Table)

## Acknowledgements

We thank Luca Ponzoni and Ivet Bahar for valuable technical and scientific discussion, Paul Welling for generously providing the anti-ROMK antisera, and members of the Brodsky and O’Donnell labs for other discussion.

## Author contributions

Conceptualization: NHN, S. Sarangi, ZWP, JLB.

Data Curation: NHN, S. Sarangi, EMM, S. Sheng.

Formal Analysis: NHN, S. Sarangi, S. Sheng, AWP, TRK.

Funding Acquisition: ZWP, JLB, TRK.

Investigation: NHN, S. Sarangi, EMM, S. Sheng, AWP.

Methodology: NHN, S. Sarangi, S. Sheng.

Project Administration: ZWP, JLB.

Software: S. Sarangi, ZWP.

Supervision: ZWP, JLB.

Writing – Original Draft Preparation: NHN, JLB, S. Sarangi, ZWP, S. Sheng.

Writing – Review & Editing: AWP, JLB, TRK.

