## Supplementary material for "Genome mining yields new disease-associated ROMK variants with distinct defects": All supplemental information (except S4 Table)

### **Supporting Information**

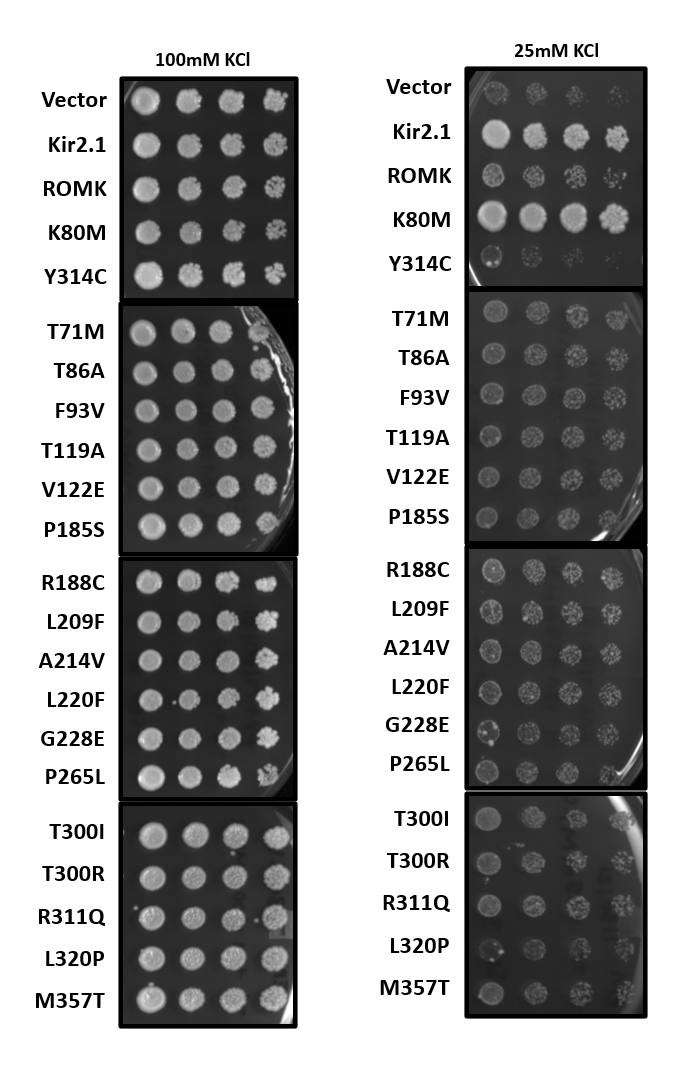

#### **S1 Figure. Viability assays of yeast expressing 17 TOPMed/ ClinVar mutations on solid medium supplemented with high (100 mM) or low potassium (25 mM).**

Representative plates from one of four independent experiments are shown. For a summary of the growth phenotypes, see **S3 Table**. Yeast cultures were transformed with an empty expression vector as a negative control, or with a plasmid expressing Kir2.1, ROMK, or the indicated ROMK mutation. Kir2.1 and a known hyperactive ROMK mutation, K80M, were used as positive controls, while a known Bartter mutation (Y314C) was used as a negative control (1, 2). Yeast cultures were grown overnight to saturation, diluted the next day, and serial 1-to-5 yeast dilutions were spotted on solid medium containing high (100 mM) or low (25 mM) potassium.
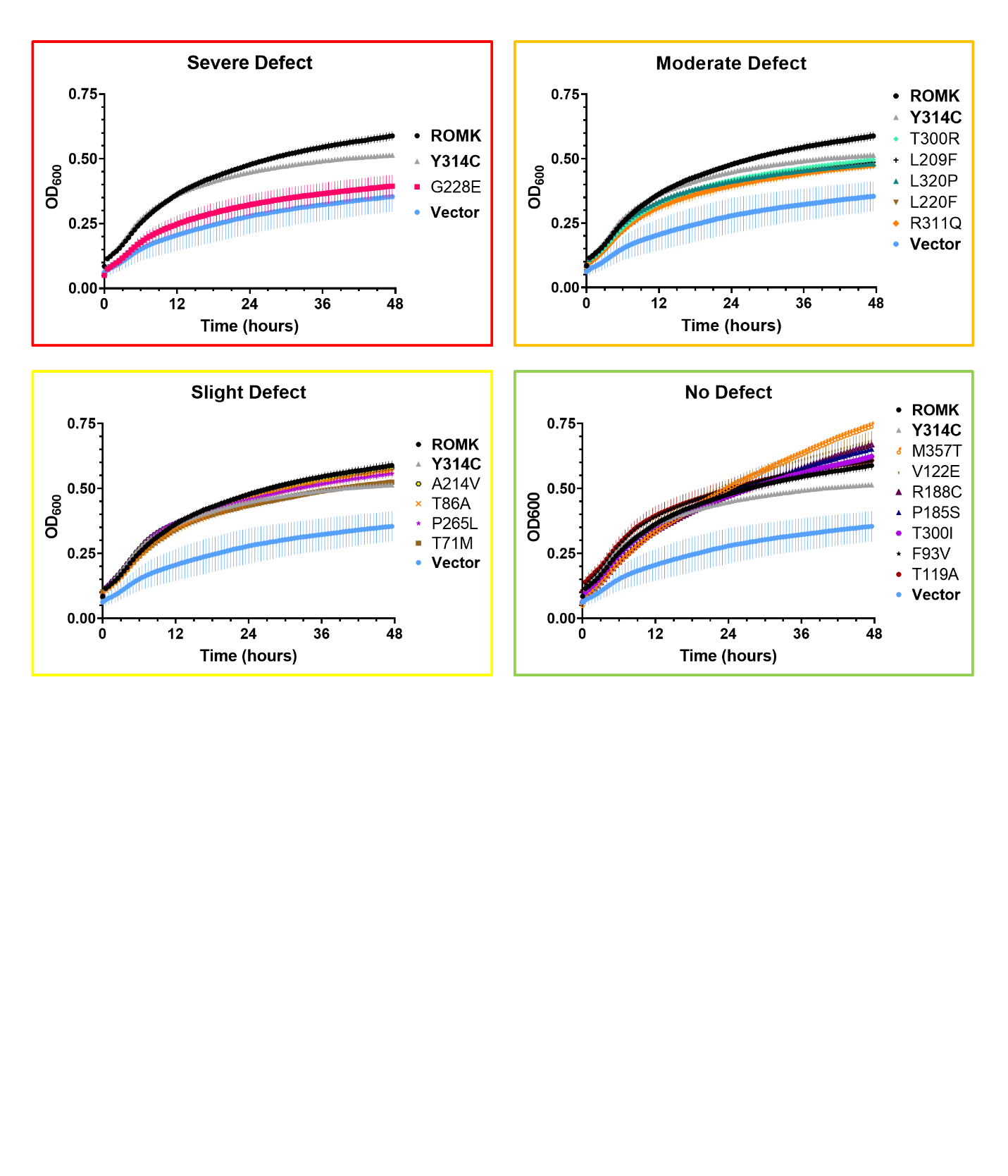

#### **S2 Figure. Viability assays of yeast expressing 17 TOPMed/ ClinVar mutations in medium containing low potassium (25 mM) grouped by growth defects.**

Yeast containing a vector control or expressing the indicated ROMK variant were grown overnight to saturation and diluted the next day to an OD_600_ of 0.20 with medium supplemented with 25 mM KCl. An empty vector and a known disease-causing ROMK mutation (Y314C) were used as negative controls (1, 2). OD_600_ readings were recorded, normalized to wells containing medium, and OD_600_ readings over the course of 48 hours were recorded. Graphs were made using GraphPad Prism (ver. 9.5.0), and error bars represent results from two replicates, ± S.E. The categorization of the relative growth defects was determined as followed: “Severe defect” for vector-like phenotype, “Moderate defect” for an intermediate phenotype between wildtype ROMK and the empty vector, “Slight defect” for a slightly-worse-than-wildtype phenotype, and “No defect” is classified as those with the same or higher growth rate as yeast expressing wildtype ROMK. For a summary of the growth phenotype categorization based on three independent experiments, see **S3 Table**.
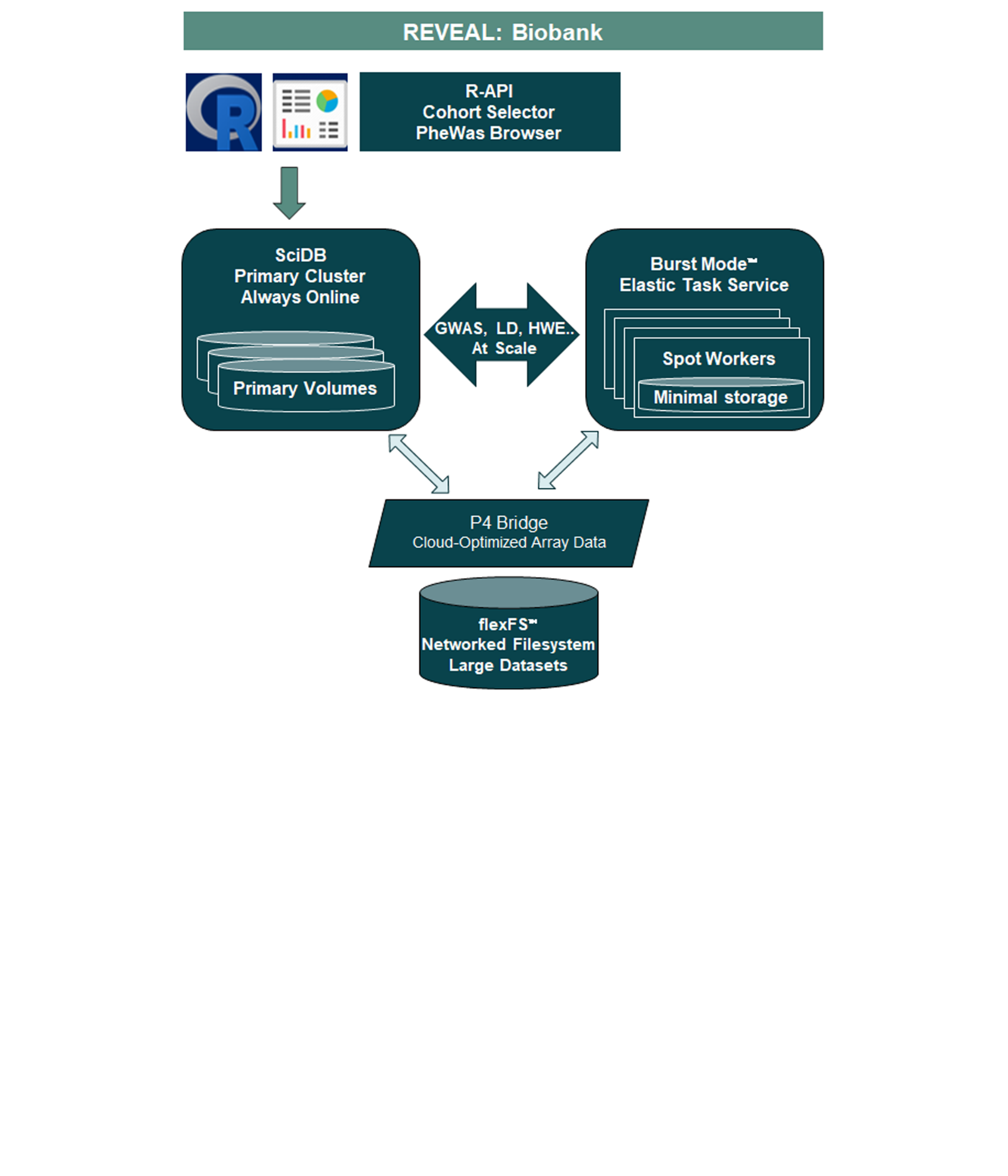

#### **S3 Figure. REVEAL: Biobank platform.**

REVEAL: Biobank is a platform comprising of an R-API, GUI’s for cohort selection & PheWas visualization, and is built upon SciDB – a computational database ideal for large scale linear algebra operations. This platform has multiple features: elastic scaling (Burst Mode) for efficient & cost-effective analyses, Bridge – a cloud-optimized array format, and flexFS – a networked POSIX compliant filesystem for working with big data in the UK Biobank. See Materials and Methods for details.

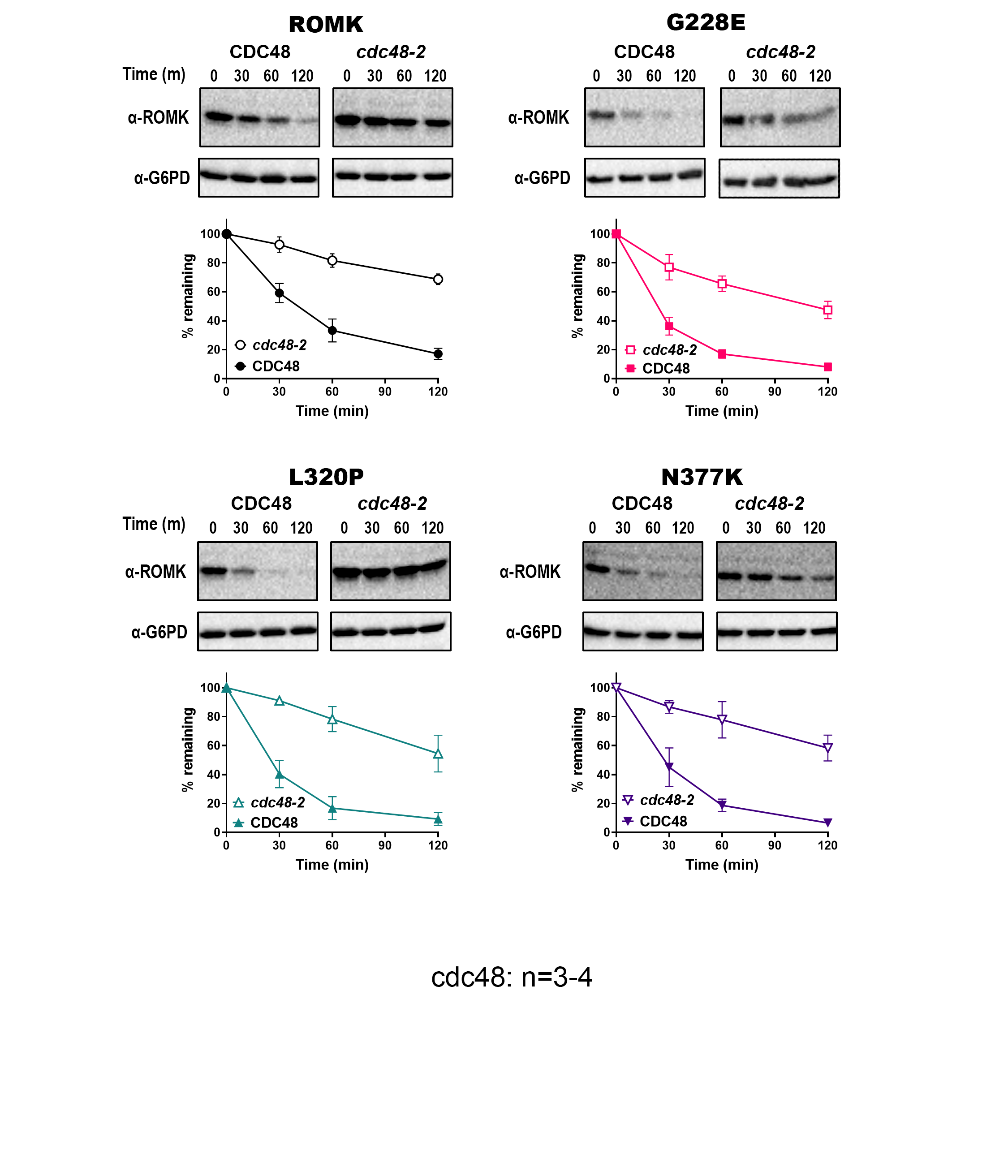

#### **S4 Figure. The degradation of select ROMK mutants is Cdc48-dependent in yeast.**

Stability assays performed in yeast expressing the wildtype ROMK, or ROMK carrying the mutation G228E, L320P, or N377K. To assess the effect of the yeast AAA^+^-ATPase Cdc48 on protein degradation, a temperature sensitive yeast strain (*cdc48-2*) was used. Yeast cultures were grown to mid-log phase (OD_600_ 0.7-1.5) at a permissive temperature, diluted, and incubated at a non-permissive temperature of 39°C for 2 hours before adding cycloheximide. Cells were then processed, and immunoblot analysis was performed (see **Materials and Methods**). A rabbit antisera was used to detect ROMK (3), and a rabbit monoclonal antibody against G6PD was used as a loading control. Representative immunoblots are shown, and graphs show the percentage of the protein remaining over time, compared to the 0-minute (m) time point, as quantified by ImageJ (ver. 1.53c). Graphs were made using GraphPad Prism (ver. 9.5.0), and data represent the means of at least three independent experiments, ± S.E. (error bars). For each experiment, a representative immunoblot is shown.

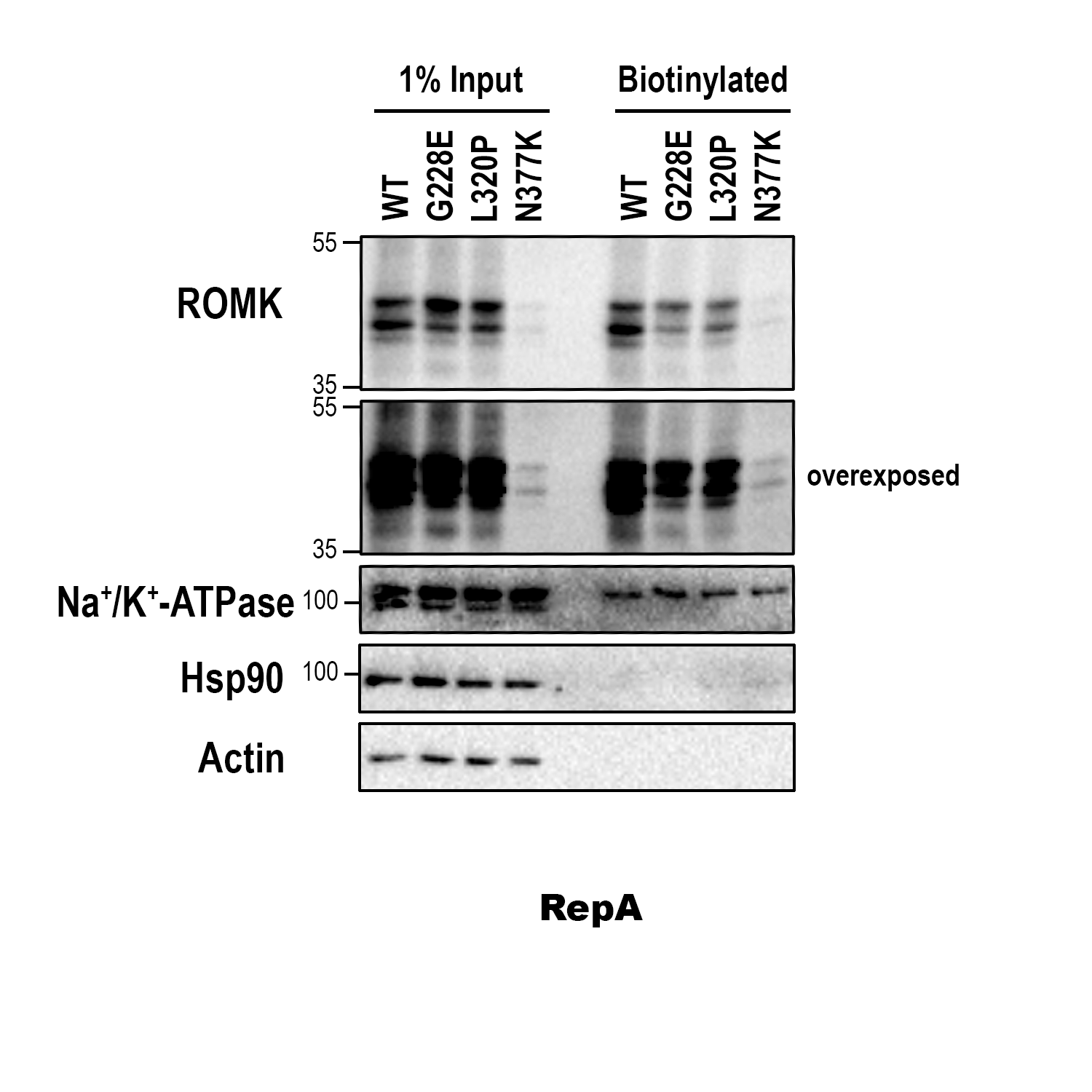

#### **S5 Figure. Select mutants reduce ROMK protein levels at the cell surface.**

Representative cell-surface biotinylation assay showing the surface expression levels of the indicated ROMK variants expressed in HEK293 cells. The experimental set-up is identical to the experiment shown in **Fig 7**, except data for N377K are included.

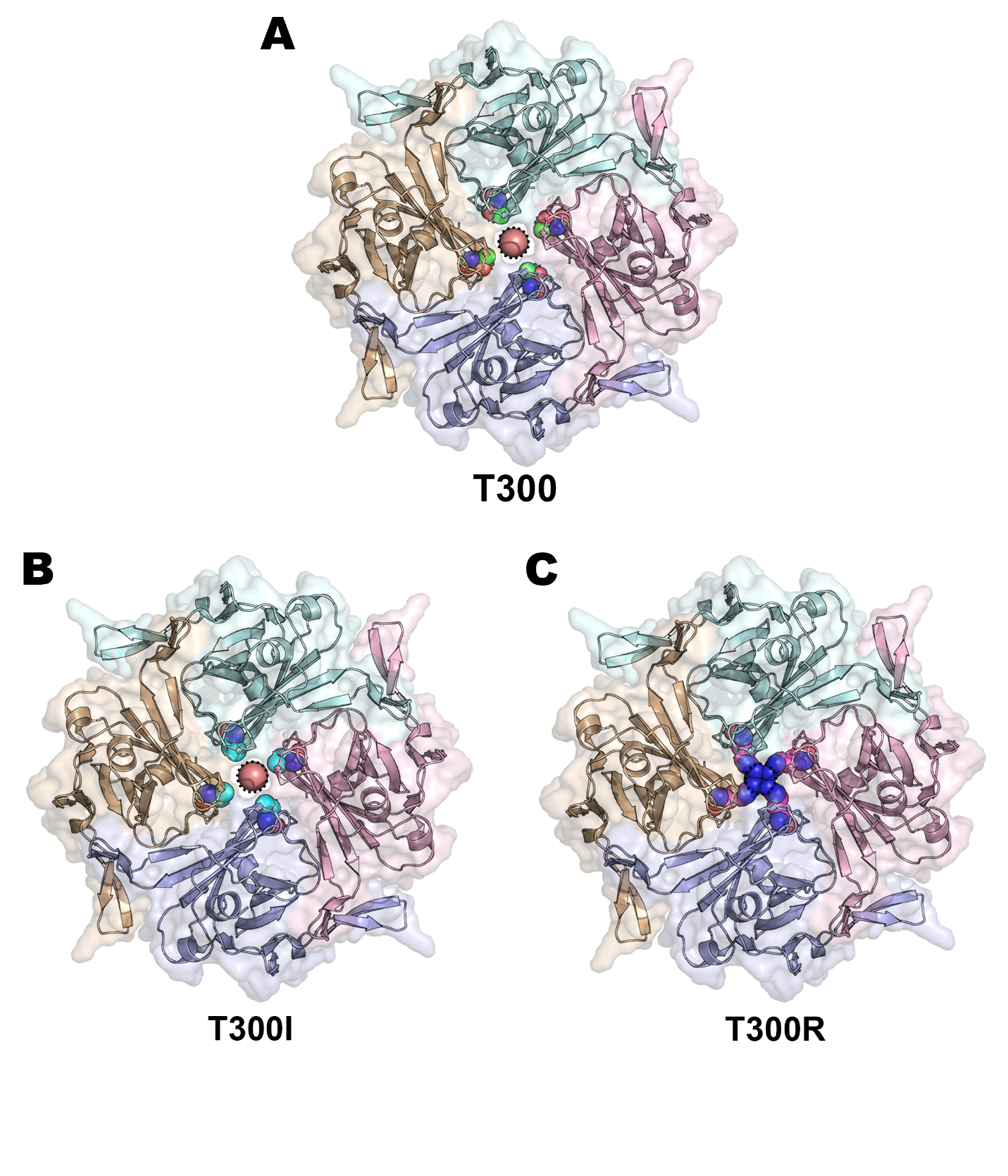

#### **S6 Figure. Structural modeling suggests the T300R mutation occludes the cytoplasmic pore.**

Homology model shows the pore of the tetrameric ROMK channel, as viewed from the cytoplasmic side. Potassium ions (spheres in the center of the pore) are shown in salmon and outlined with dotted black lines. Spheres depicting the positions of T300 **(A)**, T300I **(B)**, and T300R **(C)** are in green, cyan, and magenta, respectively. Only residues 184-364 of each chain are shown for clarity. The homology model was built based on the crystal structure of Kir2.2 (PDB ID: 3SPG), which is 47.42% identical to ROMK1. Images were rendered using PyMOL (ver. 2.6.0).

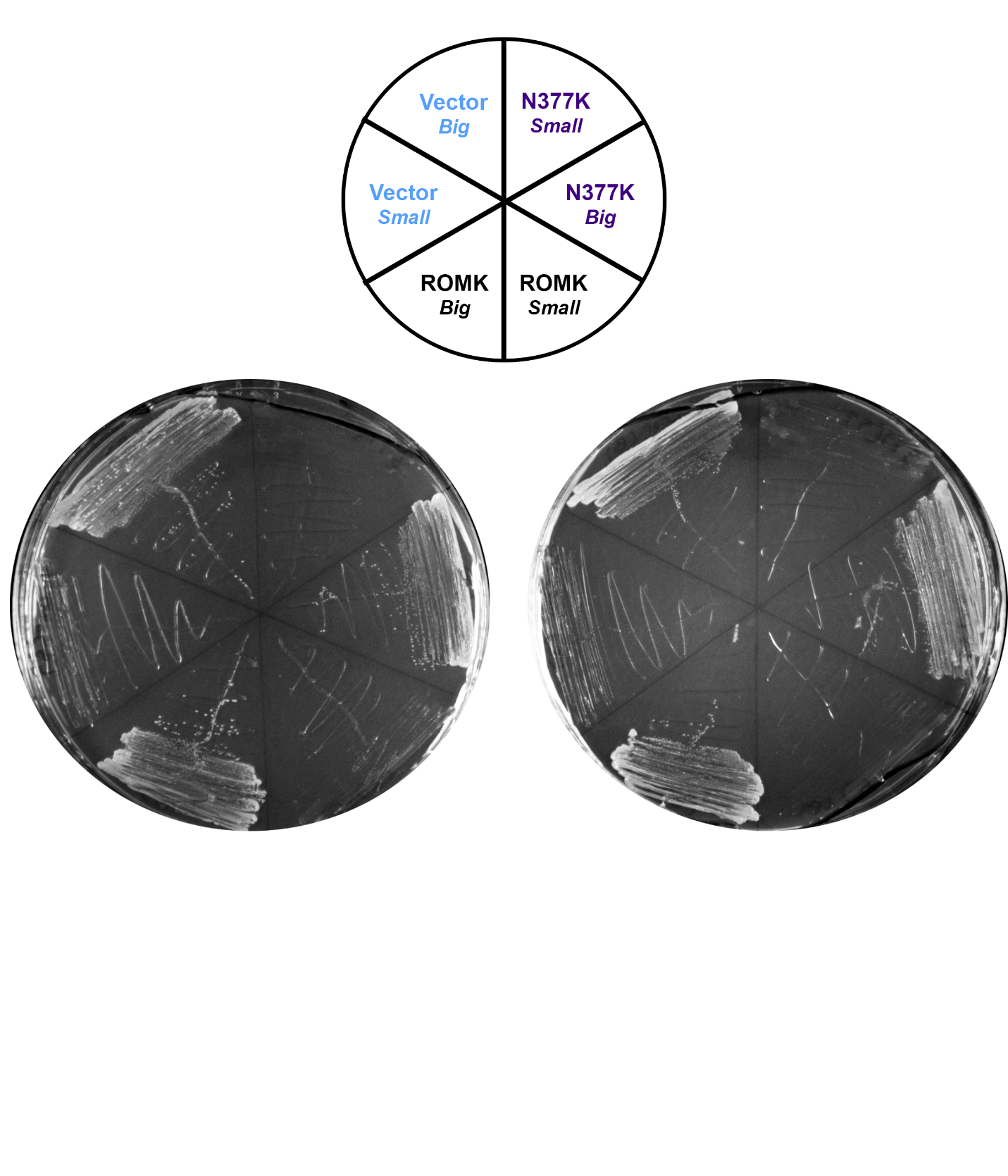

#### **S7 Figure. Slow growing *trk1*Δ*trk2*Δ yeast colonies fail to propagate on a nonfermentable carbon source.**

Large and small colonies of *trk1*Δ*trk2*Δ yeast carrying an empty vector, the ROMK protein, or the N377K mutant were propagated on synthetic complete medium lacking leucine and containing 3% glycerol. Plates show two independent replicates, and images were taken with the Bio-Rad ChemiDoc XRS+ imager after a 7-day incubation (4 days at 30°C, 3 days at 22°C).

| **Mutation** | **Rhapsody score** | **Rhapsody prediction** |
| --- | --- | --- |
| A3V | - | - |
| A6W | - | - |
| R6Q | - | - |
| T11M | - | - |
| T11A | - | - |
| T17A | - | - |
| S19N | - | - |
| R25W | - | - |
| R25Q | - | - |
| K26N | - | - |
| W27S | - | - |
| V29I | - | - |
| R31C | - | - |
| R31H | - | - |
| H35N | - | - |
| R37W | - | - |
| R37Q | - | - |
| D46N | 0.694 | Del |
| G47R | 0.905 | Del |
| C49R | 0.711 | Del |
| F53C | 0.432 | Neu |
| E57K | 0.065 | Neu |
| V66A | 0.067 | Neu |
| **T71M** | **0.790** | **Del** |
| T82I | 0.314 | Neu |
| I85N | 0.512 | Prob. Del |
| **T86A** | **0.063** | **Neu** |
| F88C | 0.739 | Del |
| **F93V** | **0.671** | **Del** |
| F94S | 0.470 | Prob. Neu |
| L97I | 0.501 | Prob. Del |
| A103V | 0.311 | Neu |
| I105V | 0.047 | Neu |
| P110L | 0.128 | Neu |
| N117S | 0.220 | Neu |
| H118Y | 0.340 | Neu |
| H118R | 0.273 | Neu |
| **T119A** | **0.298** | **Neu** |
| **V122E** | **0.796** | **Del** |
| G127S | 0.300 | Neu |
| F132L | 0.621 | Del |
| C148Y | 0.450 | Prob. Neu |
| I157N | 0.660 | Del |
| L159M | 0.613 | Del |
| S164P | 0.613 | Del |
| V168I | 0.197 | Neu |
| I170V | 0.363 | Neu |
| M174I | 0.346 | Neu |
| M174V | 0.311 | Neu |
| R184S | 0.427 | Neu |
| **P185S** | **0.549** | **Prob. Del** |
| K186N | 0.660 | Del |
| R188H | 0.710 | Del |
| **R188C** | **0.671** | **Del** |
| T193M | 0.682 | Del |
| F194S | 0.835 | Del |
| K202E | 0.443 | Neu |
| R203W | 0.285 | Neu |
| R203P | 0.598 | Del |
| R203Q | 0.543 | Prob. Del |
| G205W | 0.790 | Del |
| L207I | 0.659 | Del |
| **L209F** | **0.801** | **Del** |
| I211L | 0.453 | Prob. Neu |
| R212Q | 0.767 | Del |
| R212P | 0.850 | Del |
| **A214V** | **0.742** | **Del** |
| A214G | 0.737 | Del |
| N215D | 0.707 | Del |
| **L220F** | **0.759** | **Del** |
| **G228E** | **0.930** | **Del** |
| L230P | 0.817 | Del |
| E240Q | 0.658 | Del |
| I247V | 0.058 | Neu |
| N250S | 0.339 | Neu |
| F251C | 0.854 | Del |
| D254H | 0.597 | Del |
| A255S | 0.206 | Neu |
| A255T | 0.212 | Neu |
| N257K | 0.110 | Neu |
| **P265L** | **0.864** | **Del** |
| H270Y | 0.693 | Del |
| I272V | 0.304 | Neu |
| F278S | 0.832 | Del |
| A283V | 0.247 | Neu |
| D290G | 0.659 | Del |
| T300A | 0.737 | Del |
| **T300I** | **0.641** | **Del** |
| S305C | 0.705 | Del |
| **R311Q** | **0.729** | **Del** |
| T312S | 0.721 | Del |
| P316L | 0.824 | Del |
| E318D | 0.586 | Del |
| V319L | 0.280 | Neu |
| **L320P** | **0.566** | **Del** |
| R324C | 0.402 | Neu |
| R324G | 0.323 | Neu |
| A326S | 0.544 | Prob. Del |
| I328T | 0.636 | Del |
| S330Y | 0.820 | Del |
| R338Q | 0.316 | Neu |
| N343K | 0.699 | Del |
| K346N | 0.755 | Del |
| T347R | 0.513 | Prob. Del |
| V350M | 0.665 | Del |
| T352S | 0.602 | Del |
| M357I | 0.267 | Neu |
| **M357T** | **0.298** | **Neu** |
| L359R | 0.467 | Prob. Neu |
| N361K | 0.567 | Del |
| K363T | 0.344 | Neu |
| D364H | 0.378 | Neu |
| D364G | 0.133 | Neu |
| V365I | - | - |
| A367D | - | - |
| K370E | - | - |
| Y373C | - | - |
| N377K | - | - |
| F378L | - | - |
| I379T | - | - |
| D387H | - | - |
| D387A | - | - |
| K390T | - | - |
| M391I | - | - |

#### **S1 Table. Comprehensive list of ROMK missense mutations in the TOPMed database.**

Table shows the Rhapsody scores and predictions of 124 ROMK missense mutations from the TOPMed database that were analyzed in this study. The analysis was performed using a ROMK homology model that contains amino acids 38-364 (see **Fig 1**), so any residue outside of this range lacks a Rhapsody score, where a “-“ is shown. “Del” means a substitution is predicted to be deleterious, where “Neu” means neutral, i.e., predicted to have no effects on channel function. A designation of “Prob. Del” or “Prob. Neu” indicates that the Rhapsody score is close to the 0.5 deleterious cutoff. For example, the Rhapsody scores of I85N and F94S are 0.512 and 0.470, and thus, these mutations are categorized as “Prob. Del” and “Prob. Neu”, respectively. The mutations listed in **bold** were selected for growth analysis in yeast.

| **Mutation** | **Rhapsody score** | **Rhapsody prediction** | **Background information** |
| --- | --- | --- | --- |
| **T71M*** | 0.790 | Del | Uncharacterized Bartter |
| **T86A** | 0.063 | Neu | Bartter |
| **F93V** | 0.671 | Del |  |
| **T119A*** | 0.298 | Neu | High Frequency, uncharacterized Bartter |
| **V122E** | 0.796 | Del | Bartter |
| **P185S** | 0.549 | Prob. Del | Bartter |
| **R188C*** | 0.671 | Del | May disrupt PIP_2_-dependent gating, uncharacterized Bartter |
| **L209F** | 0.801 | Del | Bartter |
| **A214V** | 0.742 | Del | May disrupt PIP_2_-dependent gating, Bartter |
| **L220F** | 0.759 | Del | May disrupt PIP_2_-dependent gating, Bartter |
| **G228E*** | 0.930 | Del | Uncharacterized Bartter,  high Rhapsody score |
| **P265L** | 0.864 | Del | High Rhapsody score |
| **T300I*** | 0.641 | Del | G-Loop, uncharacterized Bartter |
| **T300R***^,¶^ | 0.722 | Del | ClinVar, G-Loop, uncharacterized Bartter |
| **R311Q** | 0.729 | Del | Important for inter-monomeric interactions, Bartter |
| **L320P*** | 0.566 | Del | Uncharacterized Bartter |
| **M357T** | 0.298 | Neu | Highest Frequency |

#### **S2 Table. Targeted list of 17 mutations showing Rhapsody scores, predicted phenotypes, and background information.**

Rhapsody was used to predict ROMK mutation severity based on structural, evolutionary, and dynamic features. The analysis was performed with a tetrameric ROMK homology model (Uniprot number: P48048), which was built in Swiss-Model (4) based on the crystal structure of Kir2.2 (PDB ID: 3SPG). A Rhapsody pathogenicity probability (or “Rhapsody score”) was computed for each mutation, and a “Del” (deleterious) denotation was assigned if the probability is ≥ 0.5, whereas a “Neu” (neutral) indicates a probability of < 0.5. “Prob. Del” denotes that the Rhapsody probability is close to the 0.5 deleterious cutoff (i.e., P185S probability score is 0.549). * denotes an uncharacterized Bartter mutation, which was defined as a disease-associated mutation in ClinVar, but is listed as having uncertain clinical significance. ¶ denotes the mutation obtained from ClinVar.

| Mutation | Rhapsody prediction | Growth phenotype  (Solid medium) | Growth phenotype  (Liquid medium) |
| --- | --- | --- | --- |
| ROMK | N/A | No Defect | No Defect |
| Y314C | Del | Moderate Defect | Moderate Defect |
| T71M* | Del | No Defect | Moderate Defect |
| T86A | Neu | No Defect | Slight Defect |
| F93V | Del | No Defect | No Defect |
| T119A* | Neu | No Defect | No Defect |
| V122E | Del | No Defect | No Defect |
| P185S | Prob. Del | No Defect | No Defect |
| R188C* | Del | No Defect | No Defect |
| L209F | Del | No Defect | Moderate Defect |
| A214V | Del | No Defect | Slight Defect |
| L220F | Del | Slight Defect | Moderate Defect |
| G228E* | Del | Severe Defect | Severe Defect |
| P265L | Del | Slight Defect | Slight Defect |
| T300I* | Del | No Defect | No Defect |
| T300R*^,¶^ | Del | No Defect | Moderate Defect |
| R311Q | Del | Slight Defect | Moderate Defect |
| L320P* | Del | Severe Defect | Moderate Defect |
| M357T | Neu | No Defect | No Defect |

#### **S3 Table. Table summarizing Rhapsody predictions and the growth phenotypes of the 17 TOPMed and ClinVar mutations.**

Each growth phenotype assessment was based on the results of four independent experiments for the growth assays on solid medium, and up to three independent experiments for the growth assays in liquid medium (representative growth assays shown in **Fig 2** and **S1-S2 Figs**). A known disease-causing ROMK mutation (Y314C) was used as a negative control. The categorization of growth defects was determined as followed: “Severe defect” for vector-like phenotype, “Moderate defect” for an intermediate phenotype between wildtype ROMK and the empty vector, “Slight defect” for a slightly-worse-than-wildtype phenotype, and “No defect” is classified as those with the same or higher growth rate as yeast expressing wildtype ROMK. * denotes an uncharacterized Bartter mutation, and ¶ denotes the mutation obtained from ClinVar.

#### **S4 Table. ROMK variants from the UK Biobank analyzed in this study.**

Table shows 511 *KCNJ1* variants available in the whole-exome sequencing (WES) database, which contains data from ~200k participants from the UK Biobank (5, 6). Columns represent the chromosomal location and the nucleotide change for each substitution, as well as their minor and alternative allele frequencies (denoted as maf and aaf, respectively).

*(Table submitted separately as Excel file)*

| **Phenotype** | **Chromosome position** | **Nucleotide change** | **Mutation** | **Controls** | | | **Cases** | | |
| --- | --- | --- | --- | --- | --- | --- | --- | --- | --- |
|  |  |  |  | Hom. | Het. | WT | Hom. | Het. | WT |
| Problems associated with amniotic cavity and membranes | 11:128839618 | C > T | G228E | 0 | 15 | 122037 | 0 | 2 | 532 |
| Hypertensive Heart Disease | 11:128839736 | C > T | A189T | 0 | 3 | 107271 | 0 | 1 | 167 |
| Hypopotassemia | 11:128839710 | G > A | N197N | 0 | 3 | 121124 | 0 | 1 | 531 |
| Electrolyte imbalance | 11:128839710 | G > A | N197N | 0 | 3 | 121124 | 0 | 1 | 564 |
| **Phenotype** | **Chromosome position** | **Nucleotide change** | **Mutation** | **#** | | |  | | |
|  |  |  |  | Hom. | Het. | WT |  |  |  |
| Urea | 11:128839052 | T > A | 3’ UTR | 3206 | 33538 | 93583 |  |  |  |
| Phosphate | 11:128839052 | T > A | 3’ UTR | 2950 | 30896 | 85997 |  |  |  |
| Creatinine | 11:128839856 | C > A | V149L | 0 | 2 | 32081 |  |  |  |
| Urea | 11:128839856 | C > A | V149L | 0 | 12 | 131199 |  |  |  |
| Urea | 11:128839370 | G > A | R311W | 0 | 15 | 131197 |  |  |  |
| Sodium in urine | 11:128839736 | C > T | A189T | 0 | 4 | 133636 |  |  |  |
| Creatinine (enzymatic) in urine | 11:128839618 | C > T | G228E | 0 | 23 | 133898 |  |  |  |
| Systolic blood pressure (automated) | 11:128839170 | G > T | N377K | 0 | 27 | 129973 |  |  |  |
| Systolic blood pressure (manual) | 11:128839170 | G > T | N377K | 0 | 4 | 7800 |  |  |  |

#### **S5 Table. Genotype distribution in the UK Biobank of ROMK variants with significant associations with disease phenotypes.**

The top 5 rows of the table show the population distribution of binary disease phenotypes, i.e., “Phecodes”, and the bottom 9 rows show the distribution of the metabolite disease phenotypes. “Hom.” denotes the number of individuals homozygous for the indicated mutation, “Het.” means heterozygous, and “WT” stands for wildtype, i.e., individuals without the indicated mutation. For the binary phenotypes, both the number of controls and cases are listed.

| **Primer name** | **Sequence (5’ to 3’)** |
| --- | --- |
| **T71MF** | CATCTGGACAATGGTGCTGGACC |
| **T71MR** | GGTCCAGCACCATTGTCCAGATG |
| **T86AF** | CGTGTTCATCGCAGCCTTCTTG |
| **T86AR** | CAAGAAGGCTGCGATGAACACG |
| **F93VF** | GGGGAGTTGGGTTCTCTTTGGTC |
| **F93VR** | GACCAAAGAGAACCCAACTCCCC |
| **T119AF** | CTGACAACCGCGCTCCTTGTGTG |
| **T119AR** | CACACAAGGAGCGCGGTTGTCAG |
| **V122EF** | CGCACTCCTTGTGAGGAGAACATTAATG |
| **V122ER** | CATTAATGTTCTCCTCACAAGGAGTGCG |
| **R185SF** | GATCTCTAGATCGAAAAAACGTGC |
| **R185SR** | GCACGTTTTTTCGATCTAGAGATC |
| **R188CF** | GACCCAAAAAATGTGCTAAAAC |
| **R188CR** | GTTTTAGCACATTTTTTGGGTC |
| **L209FF** | GAAGCTCTGCTTCCTCATCCG |
| **L209FR** | CGGATGAGGAAGCAGAGCTTC |
| **A214VF** | CATCCGAGTGGTAAATCTTAGGAAG |
| **A214VR** | CTTCCTAAGATTTACCACTCGGATG |
| **L220FF** | CTTAGGAAGAGCTTCCTGATTGGCAG |
| **L220FR** | CTGCCAATCAGGAAGCTCTTCCTAAG |
| **G228EF** | CAGCCACATATATGAGAAGCTTCTAAAGAC |
| **G228ER** | GTCTTTAGAAGCTTCTCATATATGTGGCTG |
| **P265LF** | CTTCATATCCCTACTGACGATC |
| **P265LR** | GATCGTCAGTAGGGATATGAAG |
| **T300IF** | CTTTTTAGATGGCATAGTGGAATCCACC |
| **T300IR** | GGTGGATTCCACTATGCCATCTAAAAAG |
| **T300RF** | CTTTTTAGATGGCAGAGTGGAATCCACC |
| **T300RR** | GGTGGATTCCACTCTGCCATCTAAAAA |
| **R311QF** | CTGCCAGGTCCAAACGTCATACG |
| **R311QR** | CGTATGACGTTTGGACCTGGCAG |
| **L320PF** | GTCCCAGAGGAGGTGCCTTGGGGCTACCGTTTC |
| **L320PR** | GAAACGGTAGCCCCAAGGCACCTCCTCTGGGAC |
| **M357TF** | CTCACTGTGCCACTTGCCTCTATAATG |
| **M357TR** | CATTATAGAGGCAAGTGGCACAGTGAG |
| **Y314C_f** | GTCCGCACGTCATGCGTCCCAGAGGAG |
| **Y314C_r** | CTCCTCTGGGACGCATGACGTGCGGAC |
| **V149L_UKBB_QC_fwd** | ATAGGTTACGGATTCAGGTTTTTGACAGAACAGTGCG |
| **V149L_UKBB_QC_rev** | CGCACTGTTCTGTCAAAAACCTGAATCCGTAACCTAT |
| **R311W_UKBB_QC_fwd** | CAACCTGCCAGGTCTGGACGTCATACGTCCC |
| **R311W_UKBB_QC_rev** | GGGACGTATGACGTCCAGACCTGGCAGGTTG |
| **A189T_UKBB_QC_fwd** | GATCTCTAGACCCAAAAAACGTACCAAAACCATTACGTTCAGCAAGA |
| **A189T_UKBB_QC_rev** | TCTTGCTGAACGTAATGGTTTTGGTACGTTTTTTGGGTCTAGAGATC |
| **N377K_UKBB_QC_fwd** | AGAGGCTATGACAACCCTAAATTTGTCTTGTCAGAAGTTG |
| **N377K_UKBB_QC_rev** | CAACTTCTGACAAGACAAATTTAGGGTTGTCATAGCCTCT |
| **Antibody** (Dilution) | **Information** |
| **ROMK** (1:1000-2000) | Rabbit antisera from the Welling lab, Baltimore, MD, USA (3). |
| **G6PD** (1:5000) | Rabbit polyclonal, from Sigma-Aldrich, St. Louis, MO, USA (A9521). |
| **Hsp90** (1:1000) | Mouse monoclonal, from Enzo Life Sciences, Farmingdale, NY, USA (ADI-SPA-830-D). |
| **Na^+^/K^+^-ATPase** (1:1000) | Mouse monoclonal, from Developmental Studies Hybridoma Bank, Iowa City, IA, USA (a5). |
| **β-Actin** (1:5000) | Mouse monoclonal, from Abcam, Cambridge, UK (ab6276). |
| **Rabbit** (1:5000) | Goat, horseradish peroxidase (HRP)-conjugated, from Cell Signaling Technology, Danvers, MA, USA (7074S). |
| **Mouse** (1:5000) | Horse, horseradish peroxidase (HRP)-conjugated, from Cell Signaling Technology, Danvers, MA, USA (7076S). |
| **Yeast strain** | **Genotype (Origin)** |
| ***trk1*Δ*trk2*Δ** | *MATα his3*Δ *leu2*Δ *ura3*Δ *trk1*Δ*::URA3 trk2*Δ*::NATMX can1*Δ*::STE2pr-HIS3* (7). |
| ***pdr5*Δ** | *MATα, his3*Δ, *leu2*Δ, *ura3*Δ, *pdr5::KANMX* (Invitrogen, Waltham, MA, USA). |
| **BY4742** | *MATα his3*Δ*, leu2*Δ*, ura3*Δ (Invitrogen, Waltham, MA, USA). |
| ***cdc48-2*** | *MATα his3*Δ, *leu2*Δ, *ura3*Δ, *cdc48-2::KANMX* (8). |

#### **S6 Table. Primers, yeast strains, and antibodies used in this study.**
